# Extracellular vesicle-mediated promotion of myogenic differentiation is dependent on dose, collection media composition, and isolation method

**DOI:** 10.1101/2022.08.22.504734

**Authors:** Britt Hanson, Mariana Conceição, Yulia Lomonsova, Imre Mäger, Pier Lorenzo Puri, Samir EL Andaloussi, Matthew J.A. Wood, Thomas C. Roberts

## Abstract

Extracellular vesicles (EVs) have been implicated in the regulation of myogenic differentiation. We observed that treatment of C2C12 murine myoblasts with either GW4869 (to inhibit exosome biogenesis) or heparin (to inhibit EV uptake) reduced myogenic differentiation. Conversely, conditioned media collected from differentiated C2C12 myotubes enhanced myogenic differentiation. Ultrafiltration-size exclusion liquid chromatography (UF-SEC) was used to isolate pure EV preparations and extracellular protein from C2C12 myoblast- and myotube-conditioned media in parallel. UF-SEC purified EVs promoted myogenic differentiation at low doses (≤2×10^8^ particles/ml), had no effect at 2×10^10^ particles/ml, and inhibited myo<genic differentiation at the highest dose tested (2×10^11^ particles/ml). Similar effects were observed with both myoblast- and myotube-derived EVs. Given that muscle-enriched miRNAs (myomiRs) are largely absent in myoblast cultures, these findings are indicative of a myomiR-independent mechanism underlying the observed pro-myogenic effects. Indeed, individual myomiRs were found to be scarce in EVs (e.g. the most abundant myomiR, miR-133a-3p, was present at 1 copy per 195 EVs). UF-SEC-purified extracellular protein had no effect on myogenic differentiation when collected in serum-free DMEM. However, a potent pro-myogenic effect was observed when Opti-MEM was used as EV harvest media. Opti-MEM contains insulin, which was sufficient to recapitulate the pro-myogenic effect. Similarly, when EVs were isolated by polymer-based precipitation, a pro-myogenic effect was observed, but only when Opti-MEM was used as a collection media. These findings highlight Opti-MEM as a potential confounding factor, and provide further evidence that polymer-based precipitation techniques should be avoided in EV research.

## Introduction

Extracellular vesicles (EVs) are nano-sized vesicles secreted by the majority of cells, and which are involved in cell-to-cell communication via the transfer of biological macromolecules (e.g. DNA, RNA and protein) [1]. Of particular interest are extracellular microRNAs (ex-miRNAs) which have been proposed to act as paracrine signalling factors. The myomiRs (miR-1, miR-133a, and miR-206) are a set of miRNAs that are highly enriched in skeletal muscle, and are involved in the regulation of myogenic differentiation [2–4]. Serum myomiRs have been proposed as minimally-invasive biomarkers in the context of multiple inherited and acquired muscle pathologies [5]. For example, myomiRs are highly elevated in the serum of Duchenne muscular dystrophy (DMD) patients and dystrophic animal models [6–12]. Release of ex-myomiRs is subject to a degree of selectivity, and is associated with muscle turnover, periods of muscle regeneration, and myogenic differentiation of myoblasts in culture [5,6,9,13]. MyomiRs are detectable in muscle-derived EVs [6,9,14], although the majority (∼99%) of ex-myomiRs are non-vesicular, and are instead most likely protected from exonucleolytic degradation through the formation of protein or lipoprotein complexes [6, 9].

We have proposed a model whereby mature muscle releases EV-encapsulated myomiRs which are taken up by immature cells in muscle (e.g. satellite cells), in order to promote their activation, and thereby supporting muscle growth and repair [15]. Consistent with this hypothesis, multiple studies have reported the cell-to-cell functional transfer of pro-myogenic factors, which could include myomiRs, between donor and recipient C2C12 murine myoblast cells [16, 17], C2C12 myotubes and neuronal cells (NSC-34) [18], activated myogenic progenitor cells (satellite cells) and myofibroblasts [19], human skeletal myoblasts and human adipose-derived stem cells [20], and human mesenchymal stem cells (MSCs) and mouse myofibres *in vivo* [21]. The transfer of EVs resulted in enhanced myogenic differentiation in recipient cell cultures [16,20,22], and improved wound repair *in vivo* [21]. Importantly, these studies have utilized methods of EV isolation that are known to co-purify other non-vesicular proteins and soluble factors, such as ultracentrifugation (UC) [20, 22] and commercial exosome isolation kits [16, 21], which could have confounded the pro-myogenic effects observed.

While cell-to-cell transfer of EV-associated miRNAs has been widely reported, the biological significance of non-vesicular ex-miRNAs, if any, is less clear. Notably, transfer of high density lipoprotein (HDL)-associated myomiRs was found to be ineffective in an *in vitro* model of cardiovascular disease [23].

Here we have investigated possible roles of vesicular and non-vesicular paracrine signalling in myoblast cultures using high purity methods of EV isolation. We show that EVs can promote myogenic differentiation, although opposite phenotypic outcomes are also observed depending on the dose used. This effect was also shown to be miRNA independent. Conversely, extracellular protein prepared in parallel with EVs did not affect myogenic differentiation unless Opti-MEM (which contains insulin) was used as an isolation medium. Similarly, the use of polymer precipitation to isolate EVs also stimulated myogenic differentiation in an Opti-MEM-dependent manner. This study has identified Opti-MEM as a potential confounding factor in EV transfer experiments and adds to a growing body of evidence that polymer precipitation techniques should be avoided in EV research.

## Materials and Methods

### Cell culture

C2C12 *Mus musculus* myoblast cells (obtained from ATCC, cat no. CRL-1772) were maintained in Growth Medium (GM: DMEM supplemented with 15% FBS and 1% Antibiotic-Antimycotic, all Thermo Fisher Scientific, Abingdon, UK). Myogenic differentiation was initiated by switching to low serum Differentiation Medium (DM: DMEM supplemented with 2% horse serum and 1% Antibiotic-Antimycotic, all Thermo Fisher Scientific).

For typical experiments, C2C12 myoblasts were seeded at a density of 1×10^5^ cells in 0.5 ml of GM per well in a 24 well multiplate, and cultured for 2 days. Subsequently, cultures were switched to DM and treated as appropriate.

EV isolation was performed on cultures grown in vesicle-free isolation medium using either DMEM supplemented with 1% Antibiotic-Antimycotic (hereafter: DMEM), or Opti-MEM I Reduced Serum Medium (Thermo Fisher Scientific, Cat # 31985062) supplemented with 1% Antibiotic-Antimycotic (hereafter: Opti-MEM), as appropriate.

### Small molecule and protein treatments

For small molecule treatments, cells were treated at the time of switching to DM with 10 µM of GW4869 (Sigma-Aldrich, Gillingham, UK), an inhibitor of neutral sphingomyelinase 2 (nSMase2) in dimethyl sulfoxide (DMSO, Sigma-Aldrich) to impair EV biogenesis [24], or 10 µM of heparin sulphate in water (Sigma-Aldrich) to prevent EV uptake or re-uptake [25]. For treatment with proteins, cells were treated at the time of switching to DM with bovine serum albumin, BSA (Thermo Fisher Scientific), insulin (from bovine pancreas, Sigma-Aldrich), or human transferrin (Sigma-Aldrich). BSA was used at a final concentration of 5 µg/ml, insulin and transferrin were used at final concentrations of 1 and 10 µg/ml, as appropriate. Untreated cultures (DM only) and/or cultures treated with DMSO were included as controls.

### RNA interference

Cells were treated with 100 µM of ON-TARGETplus Mouse *Rab27a* (11891) or *Rab27b* (80718) siRNA SMARTpools (Dharmacon, Lafayette, CO, USA) complexed with Lipofectamine RNAiMAX (Thermo Fisher Scientific) according to manufacturer’s protocols. Transfection complexes were prepared in DMEM containing no supplements (Thermo Fisher Scientific). Target regions of the ON-TARGETplus SMARTpool siRNAs are listed in **Table S1**. ON-TARGETplus Non-targeting Pool (Dharmacon) was used as a negative siRNA control.

### Immunofluorescence

Immunofluorescence (IF) was performed as described previously [26]. Briefly, cell cultures were washed with phosphate buffered saline (PBS) (Thermo Fisher Scientific), fixed with 4% paraformaldehyde (Santa Cruz Biotechnology, Dallas, TX, USA), permeabilized with 0.25% Triton X-100 (AppliChem, Darmstadt, Germany), and blocked with 5% BSA (Sigma-Aldrich, St Louis, MO, USA). Cells were subsequently incubated sequentially with primary and secondary antibodies as appropriate (**Table S2**). Nuclei were stained with Hoechst stain (i.e. Hoechst 33258 Pentahydrate (Bis-Benzimide)) (Sigma-Aldrich).

Microscopy images were acquired using an EVOS FL Cell Imaging Fluorescence Microscope (Thermo Fisher Scientific), with a 10×/0.25 objective lens. Images were captured using the inbuilt camera and subsequent image handling was performed in ImageJ.

Myogenic differentiation was quantified using three parameters. (i) Calculation of MHC+ area as a percentage of the total image area using a custom ImageJ script. This metric is convenient for the rapid processing of a large number of microscopy images as an initial screen. (ii) The myogenic index was defined as the percentage of nuclei contained within all MHC+ cells. (iii) The fusion index was defined as the percentage of nuclei contained within myotubes that contain three or more nuclei.

### EdU incorporation assay

Cell proliferation was assessed 24 hours post treatment using the Click-iT EdU Alexa Fluor 555 Imaging Kit (Thermo Fisher Scientific) according to the manufacturer’s instructions. The cultures were pulse incubated with 10 µM EdU (5-ethynyl-2′-deoxyuridine) for 1 hour in order to metabolically label newly synthesized genomic DNA in cells undergoing S-phase. Cells were subsequently fixed and permeabilised, followed by conjugation of Alexa Fluor 555-azide to the alkyne groups of the EdU nucleotides by click chemistry. The proliferation index was calculated as the percentage of Alexa Fluor 555-positive nuclei.

### RT-qPCR

Reverse transcriptase-quantitative polymerase chain reaction (RT-qPCR) was conducted following the MIQE (minimum information for publication of quantitative real-time PCR experiments) guidelines [27] where possible or appropriate. RNA extraction from cells was performed using the Maxwell RSC simplyRNA Tissue Kit (Promega, Madison, WI, USA) as according to manufacturer’s instructions. RNA extraction from EVs was performed using TRIzol LS Reagent (Thermo Fisher Scientific) as according to the manufacturer’s instructions.

RNA concentrations were quantified by absorbance at 260 nm using a NanoDrop 2000 spectrophotometer (Thermo Fisher Scientific), and cDNA generated using the High-Capacity cDNA Reverse Transcription Kit (Thermo Fisher Scientific). cDNA templates were amplified on a StepOne Plus real-time PCR Thermocycler (Applied Biosystems, Waltham, MA, USA) using Power SYBR Green Master Mix (Thermo Fisher Scientific) and gene-specific primers (**Table S3**). Duplicate technical replicates were performed per biological sample. Each reaction consisted of 2 μl of undiluted cDNA, 10 μl of 2× SYBR Green Master Mix, 1 μl of a 20× primer mix, and nuclease-free water to a final volume of 20 μl. Cycling conditions were as follows: initial denaturation for 10 minutes at 95°C, followed by 40 cycles of 15 seconds at 95°C and 1 minute at 60°C. Melt curve analysis was performed after completion of the cycling protocol (temperature range: 60 – 95°C, at a ramp rate of 0.3°C per second). A single peak confirmed an absence of primer-dimer products. A no template control (where water was substituted for the cDNA) was included for each assay run. Transcript levels were analysed using the Pfaffl method [28]. *Rab27a* and *Rab27b* transcript levels were detected using gene-specific primers and normalised to the stable murine reference gene *Rplp0*, and data were scaled such that the mean of the control group was returned to a value of 1.

### Absolute quantification miRNA RT-qPCR

miRNA RT-qPCR was performed as described in detail previously [29–31]. Briefly, RNA was extracted from 1×10^10^ EVs using TRIzol LS (Thermo Fisher Scientific) according to manufacturer’s instructions with minor modification as described previously [29]. 3 µl of 5 nM of synthetic miRNA mimic (cel-miR-39) was added to each sample at the phenolic extraction step as an exogenous spike control. 5 µl of total RNA was then reverse transcribed using a pool of miRNA-specific stem loop primers and the TaqMan MicroRNA Reverse Transcription Kit (Thermo Fisher Scientific) according to manufacturer’s instructions. 2 µl of resulting cDNA was used per subsequent qPCR reaction (20 µl total volume). qPCR amplification was performed on a StepOne Plus real-time PCR Thermocycler with TaqMan Gene Expression Master Mix (Thermo Fisher Scientific) using universal cycling conditions: 95°C for 10 min, followed by 45 cycles of 95°C for 15 seconds, 60°C for 1 minute. Details for all small RNA TaqMan assays are listed in **Table S4**. Absolute quantification was performed by comparing samples to standard curves consisting of serial dilutions of synthetic miRNA mimics for the miRNAs of interest (**Table S5**). Copy numbers determined in this manner were further scaled according to the levels of cel-miR-39 in order to account for differences in extraction efficiency between samples.

### Transmission electron microscopy

Quality and purity of EV-containing samples (isolated by different methods) were evaluated by transmission electron microscopy (TEM). Briefly, 10 µl of EV sample was applied to freshly glow discharged carbon formvar 300 mesh copper grids (Agar Scientific, London, UK) for 2 minutes. The grid was then blotted dry with filter paper and stained with 2% uranyl acetate for 10 seconds. The water droplet was then removed and the grid was air dried for 15 minutes. Grids were imaged using a FEI Tecnai 12 TEM at 120 kV using a Gatan OneView CMOS camera (Gatan, Pleasanton, CA, USA).

### Western blot

EVs and EV producer C2C12 cells were lysed in RIPA buffer (both Thermo Fisher Scientific) and protein concentrations were determined using the Micro BCA Protein Assay Kit (Thermo Fisher Scientific) as according to the manufacturer’s instructions.

Protein was added to 1× NuPAGE LDS Sample Buffer and 1× NuPAGE Reducing Agent (both Thermo Fisher Scientific) in water and incubated at 75°C for 10 minutes for protein denaturation. 10 µg of protein sample was loaded per well, and separated by SDS-PAGE using NuPAGE 4 to 12% Bis-Tris gel, 1.0 mm, Mini Protein Gels and the gel run at 100 V for 75 minutes in NuPAGE MOPS SDS Running Buffer (all Thermo Fisher Scientific). The proteins were transferred onto an Immobilon-fl polyvinylidene difluoride (PVDF) membrane (Sigma-Aldrich) at 100V for 1 hour on ice in NuPAGE Transfer Buffer (Thermo Fisher Scientific).

Membranes were blocked for 1 hour at room temperature in Intercept (PBS) Blocking Buffer (LI-COR, NE, USA). Primary and secondary antibody incubations were carried out in Intercept Blocking Buffer supplemented with 0.1% Tween-20 (Sigma-Aldrich) overnight at 4°C, or for 1 hour at room temperature respectively. Blots were washed with PBS supplemented with 0.1% Tween-20 after antibody incubations. Details of antibodies are described in **Table S2.**

### Ultrafiltration of myotube-conditioned media

C2C12 cells were seeded at a density of 4.7×10^5^ cells in 10 cm plates in 10 ml of GM and switched to DM 48 hours later. The DM was changed 72 hours later and the conditioned media (CM) collected after a further 72 hours from highly differentiated myotubes. The CM was centrifuged at 300 *g* for 5 minutes at 4°C to remove extracellular debris. The supernatant was transferred to a fresh 50 ml tube, and centrifuged at 2,000 *g* for 10 minutes at 4°C to pellet the remaining larger, non-vesicular particulate matter. This was then passed through a 0.22 µm filter. The resulting CM sample was then further fractionated by sequential ultrafiltration using molecular weight (Mr) cut-off filters as follows. 6 ml of the crude CM was passed through a 300 kDa Vivaspin concentrator (Sartorius AG, Göttingen, Germany) by centrifugation at 3,000 *g* for ∼15 minutes (or until the retentate was ∼300 µl). The EV-containing fraction (CM-EVs) was obtained from the retentate and the volume of the flow-through (FT) was increased to 3 ml in DM before filtration through a 100 kDa Amicon Ultra-15 centrifugal unit (MilliporeSigma, MA, USA) by centrifugation at 3,000 *g* for ∼10 minutes. The EV-depleted fraction containing particles and soluble factors of 100-300 kDa in size was obtained from the retentate and the volume of the FT was increased to 3 ml in DM once again before filtration through a 10 kDa Amicon Ultra-15 centrifugal unit (MilliporeSigma) by centrifugation at 3,000 *g* for ∼10 minutes. The EV-depleted fraction containing particles and soluble factors of 10-100 kDa in size (e.g. growth factors and cytokines) was obtained from the retentate. In summary, treatment fractions obtained by spin filtration included: (i) crude unfractionated CM, (ii) CM containing EVs and other particles >300 kDa in size, and CM depleted of EVs with particles that are (iii) 100-300 kDa, or (iv) 10-100 kDa, in size. Treatment of recipient C2C12 myoblasts was performed immediately following isolation of CM fractions.

### Isolation of EVs by UF-SEC

C2C12 myoblasts were seeded at a density of 4.7×10^5^ cells in ten 15 cm plates (Thermo Fisher Scientific), each containing 20 ml of GM. For the collection of MB-conditioned media, cells were cultured in GM for 4 days, and then switched to serum-free media (DMEM containing antibiotics/antimycotics, hereafter referred to as DMEM) for a further 24 hours. For the collection of MT-conditioned media, cells were differentiated for 2 days, followed by a further two days in serum-free isolation media (DMEM only).

In the cases of both MB and MT donor cultures, the ∼200 ml of pooled CM was transferred into eight 50 ml tubes (Thermo Fisher Scientific) and centrifuged at 300 *g* for 5 minutes at 4°C. The supernatant was then transferred to fresh tubes, and centrifuged again at 2,000 *g* for 10 minutes at 4°C. The supernatant was then pooled and filtered using a 0.22 µm filter unit. The resulting CM was kept at 4°C until ready for EV isolation.

UF-SEC was performed as described previously [32]. Briefly, CM ultrafiltration and concentration was carried out by tangential flow filtration (TFF) using a Vivaflow 50 R 10,000 MWCO Hydrosart membrane (Sartorius, Göttingen, Germany) and a Masterflex L/S Easy Load machine (Antylia Scientific, St. Neots, UK). Typically, ∼200 ml of conditioned media was concentrated to ∼15 ml by TFF. The volume was then further concentrated to <2 ml using a 10 kDa Amicon Ultra-15 centrifugal unit (Sigma-Aldrich).

Size exclusion liquid chromatography (SEC) was carried out using the ÄKTA Prime instrument (GE Healthcare, Pollards Wood, UK), equipped with an ultraviolet flow cell. The concentrated CM was loaded onto a column packed with Sepharose 4 Fast Flow resin (GE Healthcare, IL, USA). Eluates were measured at 280 nm absorbance to monitor the protein content of each fraction. The volume of each EV-containing fraction was set at 0.5 ml, and fractions 5-10 (EV-containing) were pooled prior to downstream analysis to obtain MB- and MT-EVs. The total volume was concentrated to ∼2 ml by spin filtration using an Amicon Ultra-2 Centrifugal Filter Unit with Ultracel-100 membrane (Sigma-Aldrich). The non-vesicular protein fraction volumes were each collected in 2 ml of PBS, and were used without further processing in downstream experiments.

### Polymer Precipitation of EVs

Conditioned media was collected from C2C12 cultures as described above and centrifuged at 300 *g* for 5 minutes at 4°C to pellet, supernatant was collected and then centrifuged at 2,000 *g* for 10 minutes at 4°C to remove any cellular debris, and then filter sterilised (0.22 µm). 200 ml of the processed CM was concentrated by TFF (10 kDa membrane) to a final volume of ∼15 ml for each sample. To precipitate EVs, 1 ml of ExoQuick-TC exosome precipitation solution (Stratech, Ely, UK) was added to 5 ml of concentrated CM, vortexed briefly, and then samples precipitated overnight at 4°C. Samples were then centrifuged at 1,500 *g* for 30 minutes at 4°C to pellet EVs. The supernatant was discarded and the pellet resuspended in sterile PBS.

### NTA

Nanoparticle tracking analysis (NTA) was performed to determine the concentration and size range of EVs obtained by UF-SEC using a NanoSight NS500 analysed using NTA 2.3 software (both Malvern Instruments, Worcestershire, UK). Measurements were analysed using the following settings: the camera level was set at 14, and the detection threshold set at 5. All other post-acquisition settings were automated. Samples were diluted 1:2,000 for count detection within the range of 2×10^8^ to 2×10^9^ particles per ml. Particles were counted and the mean of triplicate 30 second recordings was obtained using an automated script.

### Conditioned media and EV transfer treatments

For conditioned media transfer experiments, recipient cells were treated at the time of switching to DM with a 50:50 mixture of 0.5 ml conditioned media treatment fraction and 0.5 ml of fresh DM. Fresh media was added so that recipient cultures were not deprived of nutrients that may have been depleted by the metabolic activity of the producer cells, or by the sequential filtration of the CM. An equal volume of DM (1 ml) was added to cultures to be included as a DM only control.

EVs were collected by UF-SEC for MB or MT cultures, and particle counts determined by NTA. Recipient cultures were treated with EVs at the time of switching to DM, with doses ranging from 2×10^2^ to 2×10^11^ particles/ml.

Each extracellular protein fraction (#1-15) collected by UF-SEC was collected and protein concentration determined by Micro BCA assay (Thermo Fisher Scientific) as according to the manufacturer’s instructions. Recipient C2C12 cultures were treated with each fraction (final concentration of 1 µg/ml) at the time of switching to DM.

### Statistical analysis

GraphPad Prism 9 software was used for all statistical analyses. Statistical significance was determined by a Student’s *t* test for two sample comparisons, and by one-way ANOVA with Bonferroni *post hoc* test for comparisons between multiple groups.

## Results

### Inhibition of exosome biogenesis and EV uptake suppresses myoblast differentiation

C2C12 murine myoblasts undergo differentiation upon serum withdrawal and fuse to form multinucleate MTs, recapitulating the process of *in vivo* muscle regeneration. The secretome is altered during differentiation [33, 34], such that the factors released by differentiating C2C12 myotubes condition the culture media with a heterogeneous pool of biomolecules. To investigate whether EV-mediated cell-to-cell communication contributes to myogenic differentiation, C2C12 cells cultured in pro-differentiation conditions were treated with inhibitors of exosome biogenesis (i.e. GW4869, an inhibitor of neutral sphingomyelinase 2 [35]) and EV uptake (i.e. heparin [25]). C2C12 cells were treated with the inhibitors at the time of switching to differentiation media (DM) (**Figure 1A**) and myogenic differentiation assessed by immunofluorescence (IF) staining for myosin heavy chain (MHC) 2 days later (**Figure 1B**). Untreated (DM only) and DMSO-treated cultures were included as negative controls. GW4869 treatment inhibited myogenic differentiation, resulting in a ∼25% (*P*<0.01), and ∼50% (*P*<0.0001) reduction in the MHC+ area and fusion index relative to the untreated control group, respectively (**Figure 1C,E**). The myogenic index was reduced by ∼18%, although this effect did not reach statistical significance at the *P*<0.05 level (**Figure 1D**). The effect of heparin treatment similarly resulted in a reduction in the MHC+ area, in addition to the myogenic and fusion indices, by ∼35% (*P*<0.01), ∼35% (*P*<0.01) and ∼50% (*P*<0.0001) relative to the untreated control group, respectively (**Figure 1C-E**). In contrast, C2C12 myocyte proliferation was unaffected by either treatment (**Figure 1F**). Furthermore, treatment with DMSO had no effect on either myogenic differentiation or cell proliferation (**Figure 1B-F**). These results demonstrate that small molecule-mediated inhibition of exosome release and EV uptake impairs myogenic differentiation, and suppresses MT maturation. These data support the notion that C2C12 MT-derived EVs play an important pro-myogenic role through paracrine signalling during myogenic differentiation.

**Figure 1.**
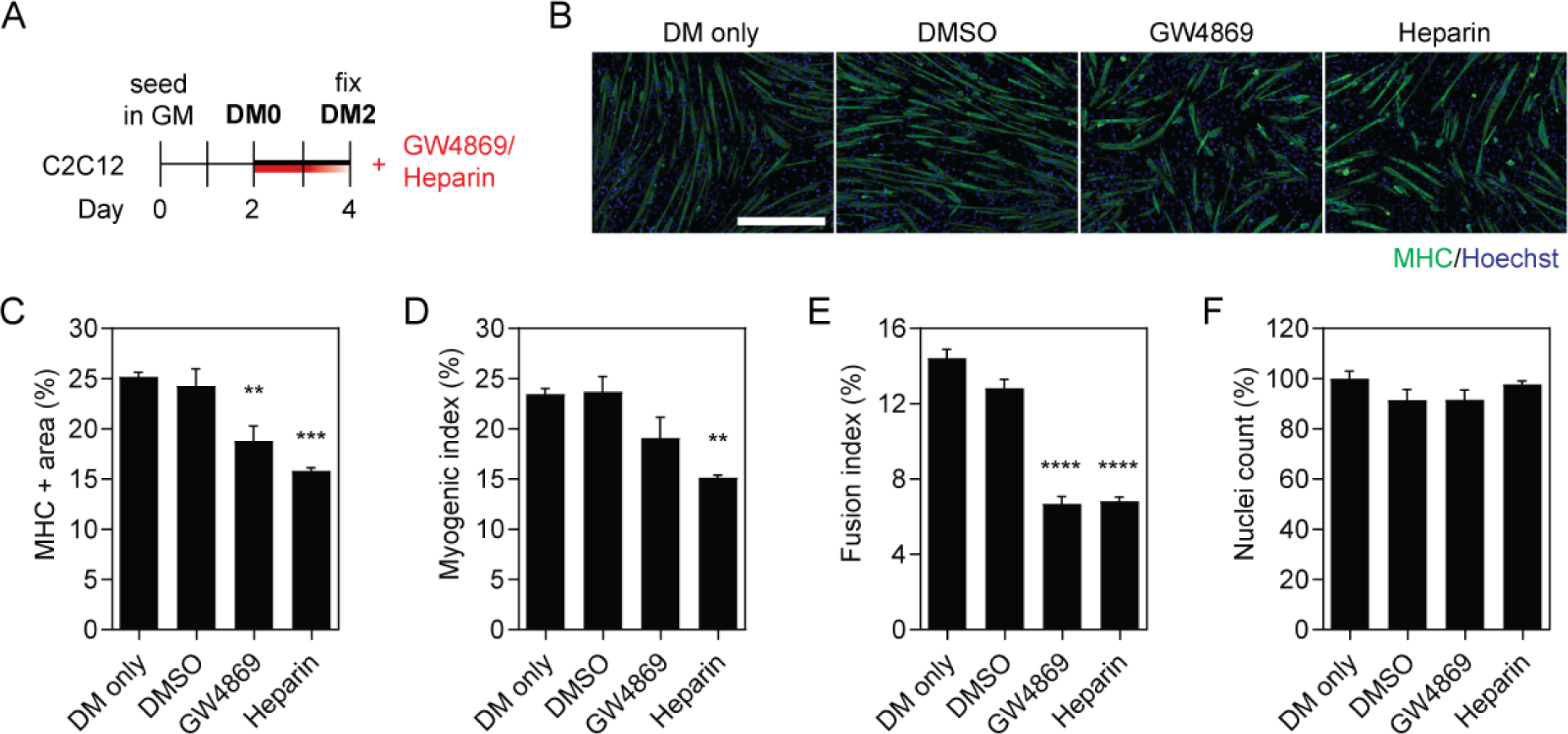
Inhibition of EV release and uptake impairs myoblast differentiation. (**A**) C2C12 cells were cultured in GM for two days and then switched to DM for a further two days. Cultures were treated with 10 µM GW4869 (exosome biogenesis inhibitor), or 10 µg/ml heparin (EV uptake inhibitor), at the time of switching to DM. Untreated (DM only) and DMSO-treated cultures were included as negative controls. (**B**) Myogenic differentiation was assessed by immunofluorescence (IF) staining for myosin heavy chain (MHC), and quantified by calculating the (**C**) MHC+ area, (**D**) myogenic index, and (**E**) fusion index. (**F**) The total number of nuclei per representative field of view are shown as a percentage relative to the control group. All microscopy images were taken at 10× magnification. Scale bar represents 400 μm. All values are mean + SEM (*n*=4). Statistical significance was determined by one-way ANOVA with Bonferroni *post hoc* test, \*\**P*<0.01, \*\*\**P*<0.001, \*\*\*\**P*<0.0001.

To assess the involvement of the specific genes associated with exosome biogenesis on myogenic differentiation, we next inhibited expression of the Rab GTPases RAB27A and RAB27B [36] by RNA interference. RAB27A and RAB27B have been found to play a role in the docking of multivesicular bodies at the plasma membrane, and knocking down these Rab GTPases decreased exosome secretion without resulting in major changes in the secretion of soluble proteins [36]. C2C12 cells were treated with short interfering RNAs (siRNAs) targeting the *Rab27a* and *Rab27b* transcripts (either separately or in combination) one day prior to switching to DM (**Figure S1A**). Transcript levels of *Rab27a* and *Rab27b* were reduced by ∼70% and ∼90% respectively, relative to the non-targeting control (**Figure S1B**). The extent of C2C12 myogenic differentiation was assessed two days later by MHC IF (**Figure S1C**). There was no significant effect on the overall MHC+ stained area between the treatments, relative to the control (**Figure S1D**). No difference in myogenic or fusion indices was observed when cultures were treated with siRNAs against either *Rab27a* or *Rab27b* alone. However, a significant >20% reduction in myogenic index (*P*<0.01) (**Figure S1E**) and >40% reduction in the fusion index (*P*<0.01) (**Figure S1F**) was observed for the combination treatment where both transcripts were knocked down simultaneously. No changes in the number of nuclei were observed for any of the siRNA treatment groups (**Figure S1G**). These results suggest that RAB27A and RAB27B are both required for the production of pro-myogenic exosomes, but that the functions of these Rab GTPases may be redundant to some extent. The level of inhibition of MT fusion with knockdown of both *Rab27a* and *Rab27b* was similar to that achieved with GW4869 or heparin treatment (**Figure 1**).

### Transfer of myotube-conditioned media and ultrafiltration-purified EVs enhances myogenic differentiation

We next sought to determine whether pro-myogenic paracrine factors secreted by C2C12 myotubes could enhance myogenic differentiation when transferred between cell culture dishes. C2C12 myotube-conditioned differentiation media (MT-CM) was harvested from cultures in the late stages of differentiation (i.e. at DM7) and then subjected to sequential ultrafiltration (UF) using 300, 100, and 10 kDa molecular weight (Mr) cut-off filters. The resulting four fractions obtained from this process therefore consisted of: (i) crude unfractionated CM, (ii) CM containing EVs and other particles >300 kDa in size, and EV-depleted CM containing soluble proteins with an Mr of (iii) 100-300 kDa, or (iv) 10-100 kDa (**Figure 2A**). Each of these fractions was mixed with an equal volume (50:50) of fresh DM and transferred to C2C12 myoblasts at DM0 (**Figure 2A**). The effect of treatment with each fraction on myogenic differentiation (by MHC IF) and cell proliferation (by EdU incorporation) was assessed in parallel experiments after 48 hours and 24 hours, respectively (**Figure 2A,B**).

**Figure 2.**
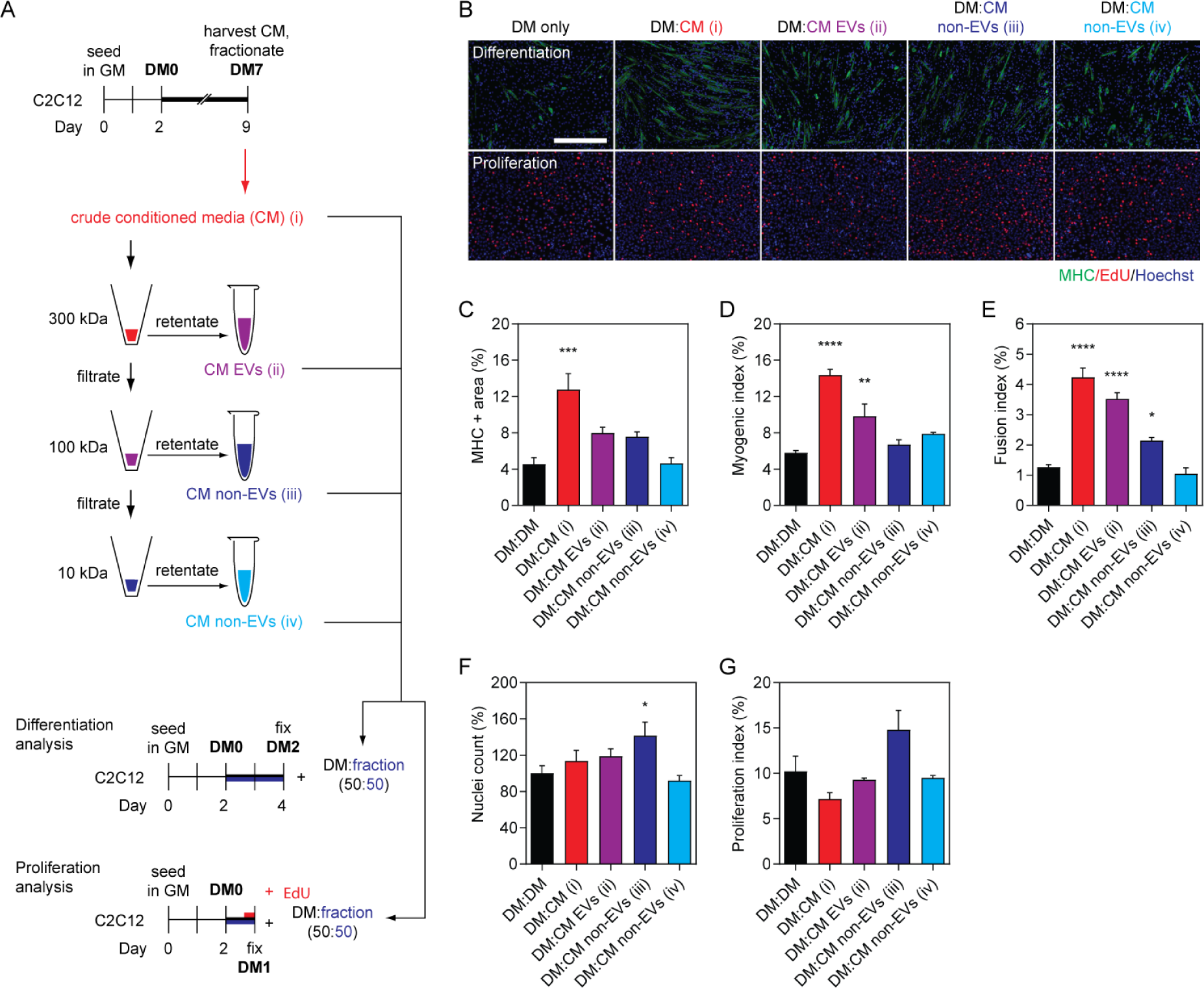
EV-containing myotube-conditioned media enhances myoblast differentiation. (**A**) C2C12 cells were cultured in GM for two days and then switched to DM for one week. Myotube-conditioned-media (CM) from DM7 cultures was fractionated using sequential molecular weight (Mr) cut-off filters of 300, 100 and 10 kDa. Treatment fractions obtained included: (i) crude CM, (ii) EV-containing CM, and (iii and iv) EV-depleted CM. Recipient cultures were treated with a 50:50 mixture of fractionated CM and fresh DM. Treatment with an equal volume of DM was included as the DM only control. Samples were collected after 48 hours for analysis of myogenic differentiation, or after 24 hours for proliferation analysis after pulsing with EdU. (**B**) Myogenic differentiation was assessed by MHC IF, and was quantified by (**C**) measuring the MHC+ area, and by calculating the (**D**) myogenic index, and (**E**) fusion index. Proliferation was assessed by determining (**F**) the percentage of nuclei per representative field of view relative to the control group, and (**G**) the proliferation index based on EdU positivity. All microscopy images were taken at 10× magnification. Scale bar represents 400 μm. All values are mean + SEM (*n*=4). Statistical significance was determined by one-way ANOVA with Bonferroni *post hoc* test, \**P*<0.05, \*\**P*<0.01, \*\*\**P*<0.001, \*\*\*\**P*<0.0001.

Treatment with crude, unfractionated MT-CM (which contains EVs as well as secreted proteins and other soluble factors) substantially enhanced myogenic differentiation (**Figure 2B**) as demonstrated by a >180% (*P*<0.001) increase in MHC+ area (**Figure 2B,C**), >145% increase in myogenic index (**Figure 2D**), as well as a >230% (*P*<0.0001) increase in the fusion index (**Figure 2E**), relative to the untreated control group. This pro-myogenic effect could not be explained by enhanced cell proliferation, as no significant difference in the total number of nuclei at DM2 (**Figure 2F**), or the proliferation index at DM1 (**Figure 2B,G**), was observed. Treatment with the EV-containing fraction (CM EVs) also enhanced myogenic differentiation (**Figure 2B**), resulting in an increase in the myogenic and fusion indices by >70% (*P*<0.01) (**Figure 2D**) and ∼180% (*P*<0.0001) (**Figure 2E**), respectively. A ∼75% increase in MHC+ area was observed which is consistent with the other two measures of myogenic differentiation for this treatment fraction, although this did not reach statistical significance (**Figure 2C**). The pro-myogenic effect of the EV-containing CM fraction could not be explained by an increase in cell proliferation (**Figure 2F,G**).

In general, treatment with either of the EV-depleted fractions had minimal effect on myogenic differentiation. A relatively small (∼70%) increase (*P*<0.05) in the fusion index was observed for the treatment group enriched in particles of 100-300 kDa (**Figure 2E**). This may be accounted for by indirect effects of increased cell density as a result of a greater number of nuclei observed only in this treatment group (*P*<0.05) (**Figure 2F**). Furthermore, the proliferation index for this group was similarly increased, although this did not reach statistical significance at the *P*<0.05 level (**Figure 2G**).

These findings suggest that differentiating MTs are capable of releasing pro-myogenic factors into the culture medium, which can be transferred to recipient cultures. Such differentiation-enhancing effects are observed for MT-derived unfractionated CM, and in the EV-containing CM fraction, but not for EV-depleted fractions.

### LC-purified EVs induce opposing effects on myogenic differentiation depending on dose

Given the pro-myogenic effects observed when MBs were treated with EV-containing MT-conditioned media, we next sought to determine if similar effects could be demonstrated for higher purity EV preparations, isolated by UF-SEC [32]. To this end, EVs were purified from C2C12 myotubes (designated as MT-EVs), and undifferentiated C2C12 myoblasts (MB-EVs, which were *a priori* not expected to promote myogenic differentiation). The resulting highly pure EV preps were transferred to separate C2C12 MB cultures at the time of initiating differentiation (i.e. DM0) (**Figure 3A**). The protein content of each fraction was measured by UV absorbance at 280 nm to monitor the liquid chromatography process, and fractions collected accordingly (**Figure 3B**). EVs elute early as a relatively well-defined peak (fractions ∼5-10), and so these fractions were collected and pooled. EV isolates were characterised by nanoparticle tracking analysis (NTA) which showed that the modal sizes for MB-EVs and MT-EVs were 108.9 nm and 83.9 nm respectively (**Figure 3C**), consistent with the expected size distributions for exosomes and other small microvesicles (i.e. ∼30-150 nm), and were not statistically different from one another (Student’s *t*-test, *P*=0.0793). Western blot analysis of EV preparations demonstrated enrichment of the EV-specific markers, Alix (PDCD6IP), TSG101, and CD9, while the cell-specific marker, Calnexin (CANX) was only detected in lysates derived from the producer cells (**Figure 3D**). Analysis of EV preparations by transmission electron microscopy revealed the presence of particles with characteristic EV-like morphology (**Figure S2**). These findings demonstrate the successful isolation of pure EVs derived from C2C12 MBs and MTs.

**Figure 3.**
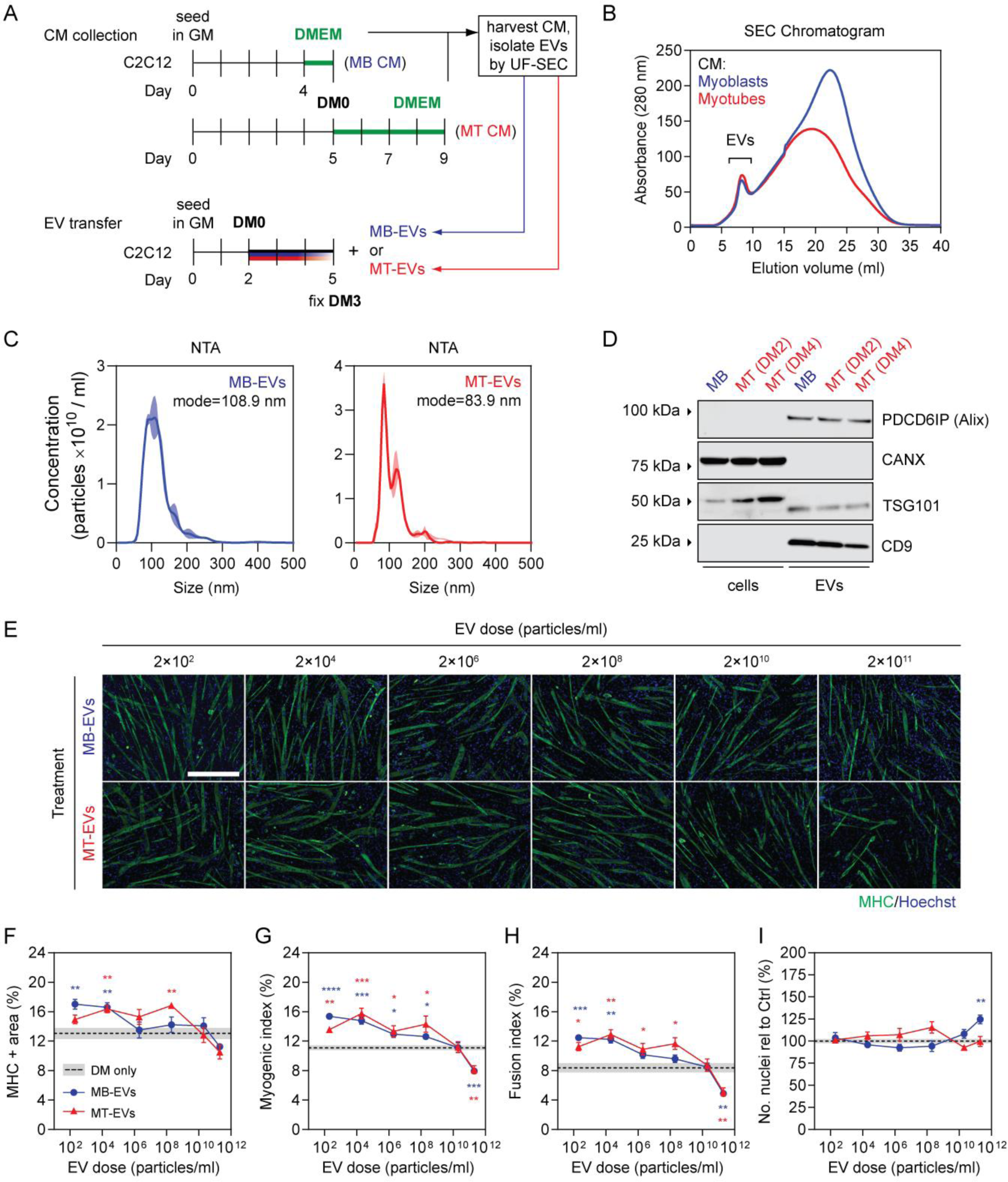
Effect of myoblast- and myotube-derived EVs on myogenic differentiation. (**A**) EVs were isolated by ultrafiltration-size exclusion liquid chromatography (UF-SEC) from CM obtained from C2C12 myoblasts (i.e. MB-EVs) or myotubes (i.e. MT-EVs). Myoblasts were grown in GM for three days and serum-free media (DMEM) for one day. Myotubes were grown in GM for five days, and then differentiation induced by serum withdrawal. The media was changed and EVs were collected in serum DMEM after a further 2 days. Recipient cultures were treated with MB-EVs or MT-EVs at doses as indicated. (**B**) The EV-containing eluates from the SEC column were determined by UV spectrophotometry absorbance at 280 nm, and pooled before treating C2C12 cells grown in GM for 2 days at the time of switching to DM. (**C**) The modal size and concentration of EVs was determined by nanoparticle tracking analysis (NTA). NTA curves represent the mean distribution from three separate isolations ± SEM (represented by the shaded area). (**D**) Exosome-specific (PDCD6IP/Alix, TSG101, and CD9) and cell-specific (CANX/Calnexin) markers were detected by Western blot. (**E**) Myogenic differentiation was assessed by MHC IF at DM3 and quantified by (**F**) measuring the MHC+ area, and using (**G**) myogenic, and (**H**) fusion indices. (**I**) The total number of nuclei per representative field of view are shown as a percentage relative to the control group. All microscopy images were taken at 10× magnification. Scale bar represents 400 μm. Values are mean ± SEM (*n*=4). Cultures treated with complete DM were used as the DM only controls. Statistical significance was determined by a Student’s *t*-test relative to the control, \**P*<0.05, \*\**P<*0.01, \*\*\**P*<0.001, \*\*\*\**P*<0.0001. Significance indicators are coloured corresponding to their respective comparisons (i.e. blue for MB-EVs vs DM only, red for MT-EVs vs DM only).

C2C12 MBs were treated according to **Figure 3A** with UF-SEC-purified EVs at doses ranging from 2×10^2^ to 2×10^11^ particles/ml, and myogenic differentiation assessed by MHC IF staining. The transfer of lower doses of MT-EVs (2×10^2^ - 2×10^8^ particles/ml) to MB cultures enhanced myogenic differentiation relative to the untreated controls (**Figure 3E-H**). The greatest effect of the MB-EV treatment group was observed at the lowest dose (2×10^2^ particles/ml), achieving a ∼30% increase (*P*<0.01) in MHC+ area (**Figure 3F**), ∼35% increase (*P*<0.0001) in myogenic index (**Figure 3G**), and ∼50% increase (*P*<0.001) in the fusion index (**Figure 3H**). Very similar effects were observed when MBs were treated with MB-EVs, especially at doses 2×10^4^ and less (Figure 3E-H). Treatment with MT-EVs resulted in the most significant promotion of myogenic differentiation at 2×10^4^ particles/ml dose, with a ∼25% increase in MHC+ area (*P*<0.01) (**Figure 3F**), a ∼40% increase in myogenic index (*P*<0.001) (**Figure 3G**), and a ∼55% increase in and fusion index (*P*<0.01) (**Figure 3H**). Conversely, at the highest dose (2×10^11^ particles/ml) both MB-EVs and MT-EVs inhibited myogenic differentiation with a ∼25-30% reduction in myogenic index (*P*<0.01) (**Figure 3G**) and a ∼40% reduction in fusion index (*P*<0.01) (**Figure 3H**). At the second highest dose tested (2×10^10^ particles/ml) there were no phenotypic effects for treatment with either MB-EVs or MT-EVs (**Figure 3E-I**).

Nuclei numbers were unaffected by treatment with either MB-EVs or MT-EVs with one exception: treatment with MB-EVs at the highest dose resulted in a ∼25% (*P*<0.01) increase in the total number of nuclei (**Figure 3I**). As such, the positive effects of MB-EV and MT-EV transfer on myogenic differentiation at all of the lower doses therefore cannot be explained by alterations in cell density.

Interestingly, the degree of enhancement observed following EV treatment (**Figure 3E-H**) was substantially lower than that observed for treatment with EV-containing CM in the previous experiment (**Figure 2B-E**). Pro-myogenic effects in the former experiments are therefore likely the result of a combination of both EV-mediated and non-EV-mediated effects.

### MyomiRs are scarce in both myoblast and myotube-derived EVs

Our lab has previously reported that myomiRs are induced by differentiation, and present at very low levels in undifferentiated MB cultures [6]. It is therefore highly unlikely that the pro-myogenic effects observed with low dose EV treatment were as a result of the functional transfer of EV-contained extracellular myomiRs, as such effects were observed with both MB- and MT-derived EVs.

To analyse the myomiR content of UF-SEC-purified EVs, we performed absolute quantification miRNA RT-qPCR on known quantities of MB-EVs and MT-EVs (determined by NTA). In this manner, the number of myomiR copies per EV could be determined. All three myomiRs (miR-1a-3p, miR-133a-3p, and miR-206-3p) were significantly upregulated (*P*<0.001) in MT-EVs relative to MB-EVs (by 13-, 28.9-, and 5.4-fold respectively) (**Figure 4**). For the MT-EVs, myomiRs were present at levels that were much lower than 1 copy per EV (miR-1a-3p: 1 copy per 5,814 EVs, miR-133a-3p: 1 copy per 195 EVs, and miR-206-3p: 1 copy per 475 EVs). These low abundance levels provide further evidence that the EV-associated pro-myogenic effect observed in C2C12 cultures is unlikely to be mediated by myomiRs.

**Figure 4.**
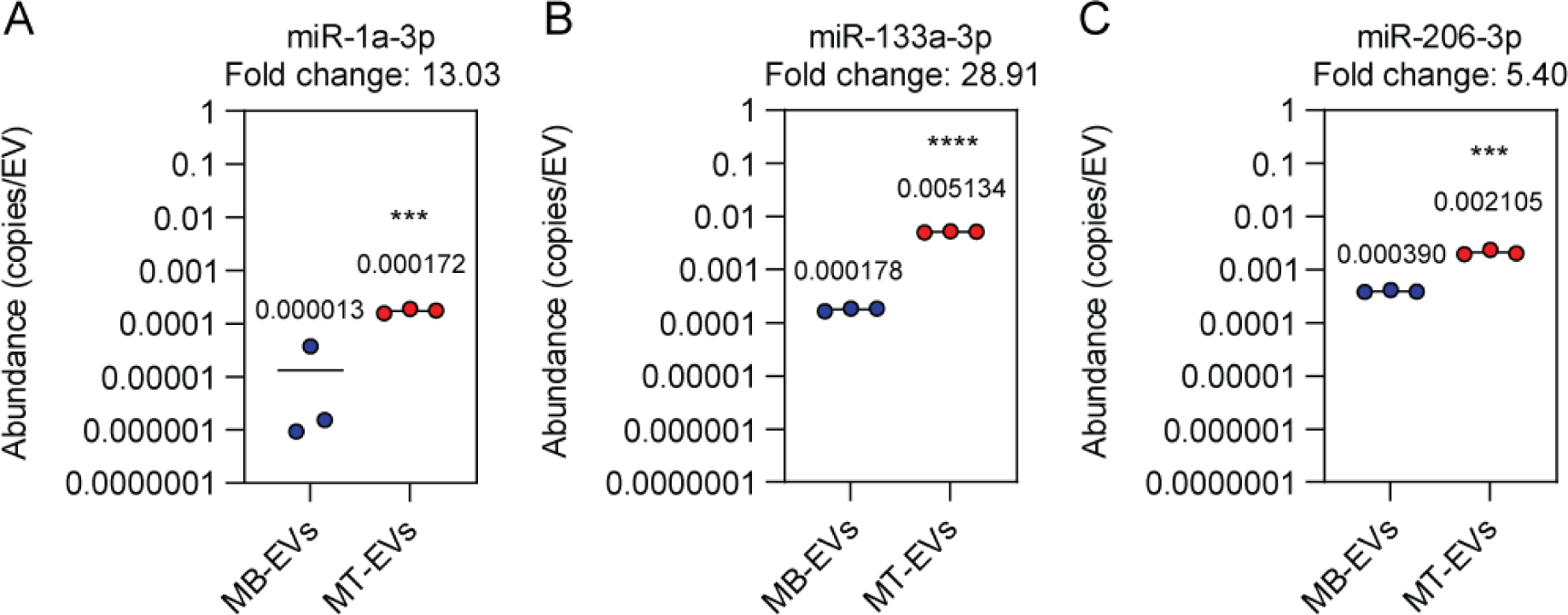
Determination of miRNA copies per vesicle in myoblast- and myotube-derived EVs. EVs were collected from myoblasts (MBs) or myotubes (MTs) as described in Figure 3A. Preparations were analysed by NTA to determine particle counts, and absolute quantification Small RNA TaqMan RT-qPCR to determine miRNA copy numbers was performed. The copies of miRNA per EV were calculated for (**A**) miR-1a-3p, (**B**) miR-133a-3p, and (**C**) miR-206-3p. Mean values are indicated (*n*=3). Statistical significance was determined by a Student’s *t*-test, \*\*\**P*<0.001, \*\*\*\**P*<0.0001

### Choice of collection medium influences pro-myogenic effects of transferred extracellular protein

We next sought to determine whether treatment with EV-depleted, soluble protein could induce pro-myogenic effects. A benefit of the UF-SEC technique is that non-EV protein-containing fractions can be collected in parallel with EV isolation using the same input CM. As such, the non-vesicular secreted protein fractions were collected from late stage (DM7) differentiating C2C12 MTs, and transferred to MB cultures at the time of switching to DM (**Figure 5A**). Fifteen protein fractions were collected based on their temporal order of elution from the column, with protein separated such that the early fractions contain higher Mr proteins, and late fractions contain lower Mr proteins (**Figure 5B**).

**Figure 5.**
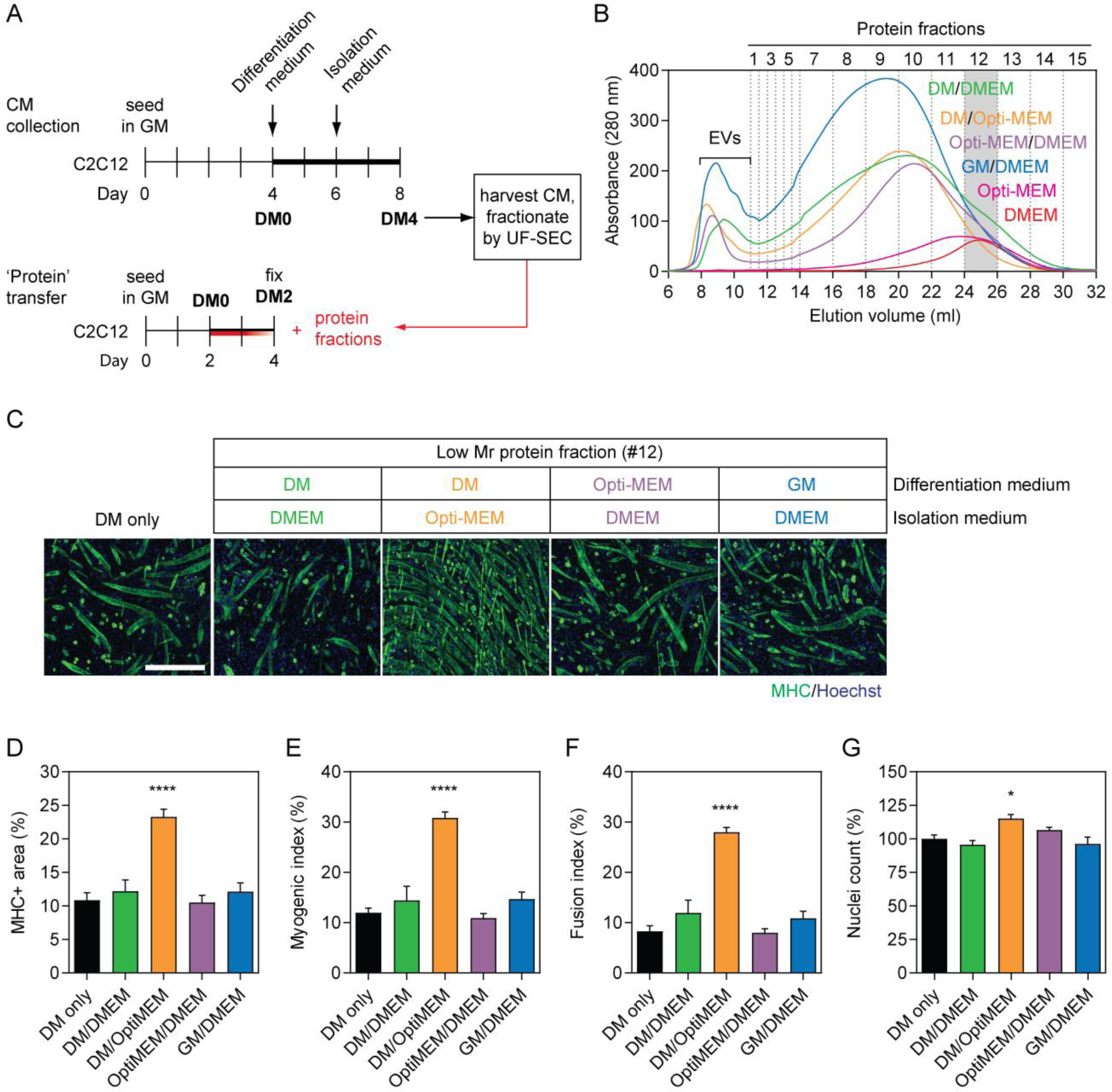
Pro-myogenic factors in Opti-MEM confound results from myotube-derived secreted protein transfer experiments. (**A**) C2C12 myotube-derived secreted protein fractions were isolated by UF-SEC from CM obtained from donor cultures with various combinations of differentiation media and isolation media: (i) DM followed by DMEM, (ii) DM followed by Opti-MEM, (iii) Opti-MEM followed by DMEM as indicated in the schematic, and (iv) GM for 3 days followed by 1 day of DMEM. (B) The protein eluates from the SEC column were measured by UV spectrophotometry absorbance at 280 nm. Equal volumes of fresh Opti-MEM or DMEM was processed by SEC in parallel. C2C12 cells were grown in GM for 2 days and then treated with (**C**) 1 μg/ml of extracellular protein #12 from each set of culture conditions at the time of switching to DM. Untreated (DM only) cultures were included as negative controls. Myogenic differentiation was assessed by MHC IF at DM2 and quantified by (**D**) MHC+ area, (**E**) myogenic index, and (**F**) fusion index. (**G**) The total number of nuclei per representative field of view are shown as a percentage relative to the control group. All microscopy images were taken at 10× magnification. Scale bar represents 400 μm. All values are mean + SEM (*n*=4). Statistical significance was determined by one-way ANOVA with Bonferroni *post hoc* test, \**P*<0.05, \*\*\*\**P*<0.0001.

Recipient C2C12 cultures were treated with 1 µg/ml of collected material from a single representative fraction (#12). We observed that there was no effect on myogenic differentiation when C2C12 cultures were treated with fraction #12 protein collected in DMEM (as described above) (**Figure 5C-F**). However, a profound increase in myogenic differentiation was observed when condition medium was collected in Opti-MEM (**Figure 5C-F**). This was represented by statistically significant (*P*<0.0001) increases in MHC+ area (>115%), myogenic index (>160%), and fusion index (>240%). The total number of nuclei were also increased in C2C12 cultures treated with protein fraction ∼12 collected in Opti-MEM by ∼15% (*P*<0.05). Such pro-myogenic or pro-proliferative effects were not observed when cells were first treated with Opti-MEM and then DMEM subsequently used as an isolation medium. This suggests that the enhanced differentiation observed above is due to co-purification of a factor from the Opti-MEM itself, rather than the result of Opti-MEM promoting the secretion of a pro-myogenic signal. Similarly, no phenotypic effects were observed when donor cells were cultured in GM, and conditioned media collected in DMEM (**Figure 5C-F**).

We next investigated whether LC fractions from unconditioned DMEM and Opti-MEM media could similarly promote myogenic differentiation. To this end, fresh DMEM and Opti-MEM were fractionated by UF-SEC and their 280 nm traces showed that they both exhibited broad peaks in the low Mr range (fractions #8-13) (**Figure 5B**). Differentiating C2C12 cultures were treated with isolated fraction #12 (1 µg/ml) collected from these fresh, unconditioned media. A strong pro-myogenic effect was observed with the protein fraction derived from Opti-MEM, but not for DMEM (**Figure S3**). Treatment of C2C12 cultures with bovine serum albumin (BSA, 5 µg/ml) had no effect on myogenic differentiation or proliferation (**Figure S4**). Taken together, these data show that protein factions collected from Opti-MEM are sufficient to recapitulate the phenomenon observed with conditioned media, and that this effect could not be explained by an increase in non-specific extracellular protein.

To investigate the Opti-MEM phenomenon further, C2C12 MBs were treated with each of the proteins fractions (collected in Opti-MEM, 1 μg/ml) at the time of switching to DM and effects on myogenic differentiation were assessed by MHC IF 24, 48, and 72 hours post treatment (**Figure S5A**). All fractions were analysed and fraction #3 and fraction #12 selected as representative of high and low Mr extracellular protein fractions, respectively (vertical grey bars on the SEC chromatogram in **Figure S5B**). The MHC+ area was progressively increased over time, and significantly increased (*P*<0.01) at 48 and 72 hours following treatment with the low Mr protein fractions in the range of fractions #7-15 (**Figure S5C,D**). This effect was greatest at 72 hours post treatment with fraction #12, resulting in a >145% increase in MHC expression relative to the control. The total number of nuclei was also significantly increased at each time point for treatment fraction #7-15, peaking with #12 at >160% relative to the DM only control (**Figure S5E**). Furthermore, cell proliferation was increased even at 24 hours following treatment (*P*<0.05) with fractions #7-15, the maximal effect being a >160% increase in the number of nuclei. Conversely, an increase in myogenic differentiation was only apparent after 48 hours post treatment. Comparatively smaller increases in myogenic differentiation (<45%) or proliferation (<30%) were observed in the high Mr protein fractions (#1-6) that elute immediately after the EV fractions relative to the DM only control, with the majority of these not reaching statistical significance (**Figure S5C-E**, significance indicators not shown for convenience). These data indicate that one or more low Mr extracellular protein induces a profound increase in myogenic differentiation and cell proliferation when Opti-MEM is used as an isolation medium. Pro-myogenic effects were observed with multiple low Mr protein fractions, suggesting that the active component protein(s) are broadly distributed across fractions. The maximal effect observed in fraction #12 corresponds with the distribution of Opti-MEM-derived proteins (**Figure 5B**). Notably, LC was performed using Sepharose 4 Fast Flow resin which has a high separation range (4×10^4^ - 3×10^7^ Da) with respect to soluble proteins, and therefore it is expected that individual proteins will be broadly distributed across multiple fractions, which was consistent with experimental observation with respect to the 280 nm SEC trace observed for fresh, unconditioned Opti-MEM (**Figure 5B**), and for the phenotypic effects observed in **Figure S5D,E**.

The exact composition of Opti-MEM culture medium is proprietary, however, according to the product datasheet (Thermo Fisher Scientific) it contains 15 µg/ml of protein in total, which includes insulin and transferrin [37]. These factors are both incidentally also components of ITS (Insulin Transferrin Selenite) medium [38], a supplement used in a number of studies to promote myogenic differentiation of cells *in vitro* [39, 40]. To test possible confounding effects mediated by these cell culture media additives, C2C12 cells were grown in GM for two days, and then treated with either insulin or transferrin (1 μg/ml) at the time of switching to DM (**Figure 6A**). The 1 µg/ml dose was selected to match the concentration of total extracellular protein transferred in the experiments described above (**Figures 5, S5**). Myoblast cultures treated with insulin exhibited a pronounced enhancement in myogenic differentiation with increases in MHC+ area expression (>95%), myogenic index (>150%), and fusion index (>230%) at DM2 (**Figure 6B-E**). Conversely, treatment with transferrin had no effect on myogenic differentiation. Assessment of the number of nuclei per well revealed that treatment with insulin resulted in a ∼150% increase (*P*<0.0001) relative to the DM only control (**Figure 6F**).

**Figure 6.**
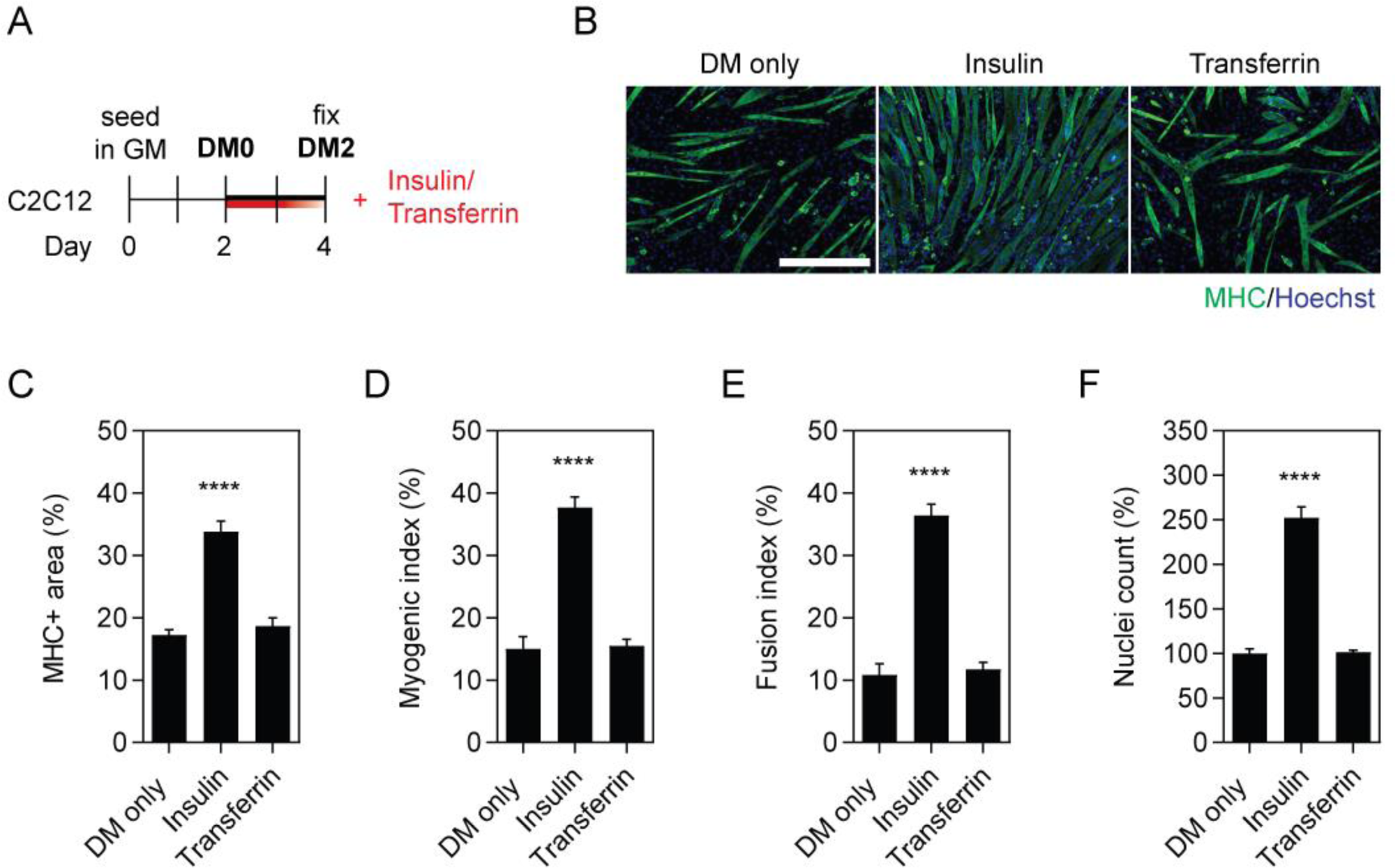
Insulin, but not transferrin, enhances myogenic differentiation in C2C12 cell cultures. (**A**) C2C12 myoblasts were grown in GM for two days, and treated with insulin or transferrin (1 or 10 μg/ml) at the time of switching to DM for two days. Myogenic differentiation was assessed by MHC IF at DM2 (**B**), and quantified by measuring the (**C**) MHC+ area, (**D**) myogenic index, and (**E**) fusion index. Untreated (DM only) cultures were included as negative controls. (**F**) The total number of nuclei per representative field of view are shown as a percentage relative to the control group. All microscopy images were taken at 10× magnification. Scale bar represents 400 μm. Values are mean + SEM (*n*=4). Statistical significance was determined by a one-way ANOVA with Bonferroni *post hoc* test, \*\*\*\**P*<0.0001.

### Vesicle-mediated pro-myogenic effects are influenced by EV isolation method

Given that we observed a confounding pro-myogenic effect of low Mr extracellular protein fractions when using Opti-MEM as a collection medium (**Figure 5**), we next sought to investigate whether the choice of EV isolation method contributes towards this phenomenon.

As such, conditioned media was collected from late stage differentiating MT cultures (DM4) in either Opti-MEM or DMEM, and then EVs isolated using either UF-SEC (designated as LC MT-EVs) or polymer precipitation (using the commercially available ExoQuick kit, designated as EQ MT-EVs) (**Figure 7A**). The protein content of each fraction obtained by UF-SEC was measured by UV absorbance at 280 nm (**Figure 5B**). EV samples from both isolation methods were characterised by NTA, indicating that the particle modal sizes were ∼71 nm and ∼97 nm for LC MT-EVs and EQ MT-EVs, respectively (**Figure 7B**), consistent with the expected sizes of exosomes and other small microvesicles. EQ MT-EVs were significantly larger than the corresponding LC EV treatment group (i.e. for both DMEM and Opti-MEM collection media) (Student’s *t*-test, both *P*<0.05).

**Figure 7.**
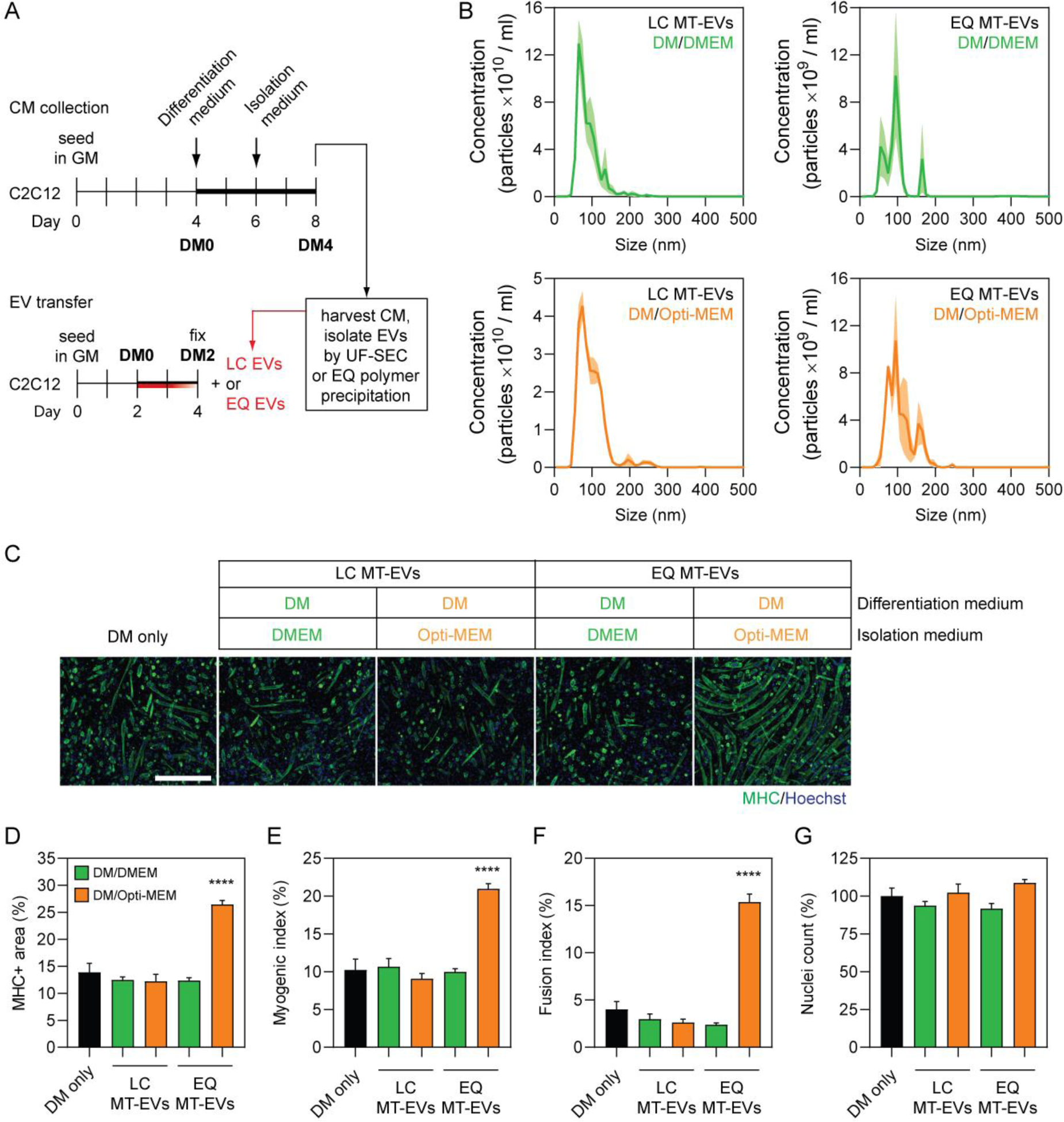
EVs obtained from Opti-MEM collection media exhibit substantial pro-myogenic effects only when using polymer precipitation isolation methods. (**A**) EVs were isolated by UF-SEC (LC MT-EVs) or ExoQuick polymer precipitation (EQ MT-EVs) from C2C12 myotube-derived CM obtained from cell growth in different isolation media. Myoblasts were grown in GM for four days, DM for two days, and isolation medium (Opti-MEM or DMEM) for a further two days. Recipient C2C12 myoblast cultures were treated with 2×10^9^ EVs/ml at the time of switching to DM. (**B**) The modal size and concentration of EVs was determined by NTA. Values are the mean distribution from three separate experiments, shown ± SEM (the shaded area). (**C**) Myogenic differentiation was assessed in recipient cultures by MHC IF at DM2 and quantified by (**D**) measuring the MHC+ area, and using (**E**) myogenic, and (**F**) fusion indices. (**G**) The total number of nuclei per representative field of view are shown as a percentage relative to the control group. Untreated (DM only) cultures were included as negative controls. All microscopy images were taken at 10× magnification. Scale bar represents 400 μm. All values are mean + SEM (*n*=4). Statistical significance was determined by a one-way ANOVA with Bonferroni *post hoc* test, \*\*\*\**P*<0.0001.

LC MT-EVs and EQ MT-EVs were transferred to MB cultures at the time of initiating differentiation, and myogenic differentiation assessed after two days (**Figure 7C**). In order to test the potential confounding effects of Opti-MEM we selected a dose of 2×10^9^ particles/ml, as this was expected to have no overall effect on myogenic differentiation as mediated by the EVs *per se*, based on our previous findings (**Figure 3E-H**). Treatment with EQ MT-EVs at this dose exhibited a highly pro-myogenic effect when isolated in the presence of Opti-MEM, with a >90% increase in both the MHC+ area and myogenic index relative to the DM only control (**Figure 7D,E**), and a >280% increase in the fusion index (all *P*<0.0001) (**Figure 7F**). Conversely, LC MT-EVs had no effect on myogenic differentiation, as expected. EVs collected in DMEM had exhibited no significant effects on myogenic differentiation when purified by either method. The number of nuclei was not altered relative to the control for any of the treatment groups (**Figure 7G**).

This pro-myogenic effect was found to be dose-dependent, with EQ MT-EV doses ≥2×10^7^ particles/ml promoting enhanced differentiation (**Figure S6A,B**), in contrast to the results observed for LC MT-EVs (**Figure 3**). Taken together, these data strongly suggest that contaminating proteins (e.g. insulin) enhance myogenic differentiation in polymer precipitated EV isolates when collected in Opti-MEM isolation media.

## Discussion

Here we have investigated the cell-to-cell signalling potential of EVs and non-vesicular extracellular protein in the context of myogenic differentiation. This study adds to the growing literature that supports the notion that EVs can mediate a pro-myogenic effect, at least to some extent. We observed that inhibition of EV release using GW4869 (a small molecule inhibitor of nSMase2 that inhibits the ceramide-dependent exosome biogenesis pathway [41]) resulted in suppression of myogenic differentiation (**Figure 1**). Additionally, siRNA-mediated knock down of *Rab27a* and *Rab27b*, two RAB GTPases that are involved in the ESCRT-dependent mode of exosome release [36,42–44], recapitulates this effect (**Figure S1**). Similarly, inhibition of EV uptake via treatment with heparin also inhibited myogenic differentiation (**Figure 1**). Conversely, myotube-conditioned media, and conditioned media that had been filtered to retain EVs, promoted myogenic differentiation in recipient C2C12 cultures (**Figure 2**). Such conditioned media preparations contain a complex mixture of biomolecules and so pro-myogenic effects could not be unambiguously attributed to EVs for this experiment. To further investigate the role of EVs specifically, we utilised UF-SEC to obtain highly pure EV preparations (**Figure 3**). Treatment of myoblast cultures with UF-SEC-purified EVs enhanced myogenic differentiation, but only at concentrations of 2×10^8^ particles/ml and below. Conversely, differentiation was inhibited at the high dose (2×10^11^ particles/ml, corresponding to ∼1 mg/ml of protein). These surprising findings highlight the importance of determining appropriate dose windows as opposite phenotypic outcomes were observed in this biological context depending on the dose. Notably, it has recently been reported that treatment with high concentrations of exogenous EVs can inhibit endogenous EV production [45].

Determining physiologically-relevant EV doses remains a significant challenge. Forterre and colleagues conducted proteomic analysis of EVs produced by C2C12 myoblasts and myotubes, and reported that differentiating C2C12 cells release 0.42 ± 0.01 µg/ml of EVs per 24 hour period [16]. This value was therefore considered to be an approximate ‘physiological dose’ for C2C12 cells in the present study. Many of the studies that have assessed the functional transfer of EVs in culture used protein concentration as a measure of dose, and phenotypic effects have been reported with 10-200 µg/ml of exogenously-derived EVs [1,20,46]. Importantly, many of the publications describing the functional role of EVs in myogenic differentiation [13,16,19– 21,47,48] have utilised commercially-available polymer precipitation kits [20] but are known to co-purify non-vesicular proteins and soluble factors [49]. Ultracentrifugation is generally regarded as the gold standard method of EV isolation and has also been used to purify myocyte-derived EVs [1, 46], but presents its own technical issues. Specifically, ultracentrifugation results in inconsistent yields and low purity as a result of co-sedimentation of non-vesicular soluble proteins and RNAs with EVs [32,60,61]. As a result, the use of protein concentration as a measure of dose may be misleading when comparing between studies that utilise distinct EV isolation methods. With this limitation in mind, the present study used MB-EVs and MT-EVs ranging from 2×10^2^ to 2×10^11^ particles/ml (corresponding to 1 pg/ml to 1 mg/ml of protein).

Our group [6, 50], and others [19, 21], have hypothesised that the protection of ex-myomiRs within EVs may enable their cell-to-cell transfer in a paracrine manner. However, in this study we observed pro-myogenic effects using EVs harvested from both differentiated and undifferentiated donor cultures. Given that myomiR levels are lowly expressed in MB donor cultures [6], this finding strongly argues against a myomiR-related mechanism for the observed pro-myogenic effects. Furthermore, EV myomiRs were similarly found to be present at very low levels (<1 copy per 195 EVs for miR-133a-3p in MT-EVs) (**Figure 4**), consistent with reports of EV miRNA concentrations in other biological contexts [51, 52]. Similarly, we have previously reported that the majority (∼99%) of extracellular myomiRs are non-vesicular in serum derived from the dystrophin-deficient *mdx* mouse [6,9,14], and are instead stabilised in soluble protein complexes. We were therefore motivated to determine if transfer of the non-vesicular majority of myomiRs could induce changes in C2C12 phenotypes. To this end, we collected non-vesicular protein fractions by UF-SEC in parallel with EV isolation. When the protein fractions were collected in serum-free DMEM media, no pro-myogenic effects were observed (**Figure 5**). These data suggest that extracellular protein-associated myomiRs cannot promote myogenic differentiation in this model system. However, when Opti-MEM was utilised as the isolation medium, the resulting protein fractions greatly enhanced myogenic differentiation (**Figure 5, S5**). Protein purified from fresh Opti-MEM and treatment with insulin (a major component of Opti-MEM) were sufficient to recapitulate this effect (**Figure 6, S3**). Together these data strongly suggest that the use of Opti-MEM as an isolation medium is a major potential confounding factor in studies of conditioned media transfer in the context of myogenic differentiation (or any other insulin-sensitive biological system).

The use of polymer precipitation methods makes it difficult to conclusively attribute a functional effect observed to EV-specific paracrine signalling, rather than some other extracellular contaminant. Indeed, an International Society for Extracellular Vesicles (ISEV) position paper has specifically highlighted the problematic nature of polymer precipitation methods in EV research as these kits ‘do not exclusively isolate EVs, and are likely to co-isolate other molecules, including miRNA-protein complexes [53]. For instance, polymer precipitation methods of EV purification have been shown to co-purify vesicle-free miRNAs from rat plasma [54], and non-vesicular extracellular Argonaute-2 (AGO2) complexes from the secretome of MCF-7 breast cancer cell cultures [55]. Furthermore, it has been suggested that the use of EV precipitation techniques has led to the misattribution of pro-angiogenic and regenerative cell signalling effects as a result of the co-purification of non-vesicular soluble factors expressed by mesenchymal stem cells [56]. The data presented in this study provide evidence to support this notion, as polymer precipitated-EVs exhibited a pro-myogenic effect only when Opti-MEM was used as the collection media (**Figure 7**). In contrast, LC-purified EVs collected in Opti-MEM did not induce similar pro-myogenic effects. These findings suggest that polymer precipitation methods co-purify media contaminants with the potential to elicit profound confounding effects. However, it has recently been reported that a protein corona forms around EVs in the circulation [57]. It is therefore possible that techniques such as UF-SEC may strip away corona proteins that are important for the phenotypic effects mediated by EVs. However, it is doubtful that exogenous insulin at supraphysiological concentrations would be considered a *bona fide* EV corona protein, and so such an eventually is unlikely to explain the results presented here.

The endocrine system plays an important role in skeletal muscle metabolism through the targeted action of receptor-mediated signalling by growth factors (e.g. insulin-like growth factors 1 and 2, IGF1 and IGF2), and hormones (e.g. growth hormone, insulin, androgens) [58]. Insulin is a low molecular weight protein (5.8 kDa) that plays an important role in skeletal muscle function [59] via intracellular signalling cascades involving the PI3K/Akt and Shc/Ras/MAP kinase axes [60]. Such pathway activation has also been reported following insulin treatment in C2C12 myoblast cultures [61, 62]. Notably, the dose of insulin used in the present study (i.e. at micromolar concentrations) is much higher than the range of circulating insulin in the body (i.e. at picomolar concentrations) [63].

While polymer precipitation methods may confound the biological interpretation of EV transfer experiments through the co-precipitation of media-derived contaminants like insulin, they may equally concentrate endogenously-derived factors. For example, a study by Hu *et al*. reported that IL6 (a soluble cytokine that signals via interaction with its cognate receptor at the membrane of recipient cells) is packaged within EVs that are then capable of cell-to-cell communication in C2C12 myotubes and 3T3-L1 adipocytes based on polymer precipitation experiments [64]. The authors own data does not support this conclusion, given that the signalling effect was blocked by anti-IL6 antibodies. Instead, it is much more likely that soluble IL6 was co-purified together with the vesicles, and the reported biological effect have been misattributed to the EVs themselves. Similarly, a study by Hettinger *et al*. used polymer precipitation to purify EVs from primary human myoblasts after the induction of senescence via treatment with hydrogen peroxide to suggest that EVs are senescence-associated secretory phenotype (SASP) mediators [65]. The SASP is a well-described effect associated with the release of multiple soluble extracellular factors (e.g. cytokines, proteases, growth factors etc.) form senescent cells [66], proteins which are likely to be co-purified in polymer precipitated EV isolates.

The data presented here add to the growing body of literature suggesting that EVs constitute a secretory signalling function in skeletal muscle, although with several caveats. Forterre *et al*. reported that MT-EVs are capable of transferring exogenous small RNAs (siRNA and non-mammalian miRNA) to myoblasts, and suggested that EV-mediated transfer of miRNA-133a-3p can suppress *Sirt1* expression in recipient cultures [48]. In a follow-up study, the same group reported that MT-EVs (but not MB-EVs) suppressed myoblast proliferation [16]. This phenotypic finding was not observed in our data, although our experiments were not primarily designed to test this directly. Both of these studies utilised ultracentrifugation for EV isolation.

Several studies have reported pro-myogenic effects of EVs *in vivo*. Nakamura *et al*. reported that mesenchymal stem cell (MSC)-derived EVs (isolated by ultracentrifugation) enhanced C2C12 differentiation, and improved regeneration after cardiotoxin-induced muscle injury [21]. Similarly, Choi *et al*. reported that EVs (polymer precipitated from media containing 10 µg/ml insulin) derived from differentiating human skeletal myoblasts could promote myogenic differentiation in human adipose-derived stem cells and reduced fibrosis in a laceration muscle injury model [20]. Multiple other studies have utilised polymer precipitation in the context of EV-mediated cell-to-cell signalling and myomiR transfer *in vivo* [19, 47].

In conclusion, this study demonstrates that highly pure preparations of EVs derived from differentiating muscle cells have the potential to transfer pro-myogenic signals to myoblasts when provided at appropriate concentrations. Notably, opposite phenotypic outcomes were observed at the highest dose of EVs administered, underlining the importance of dose response experiments in EV research. Conversely, we observed that non-vesicular protein fractions were unable to transfer pro-myogenic signals to recipient cells, provided that specific protein contaminants are absent. These findings are important as there is a risk that biological effects could be misattributed to EVs when they are in reality the result of exogenous or endogenous non-vesicular factors that are co-purified when using certain methods of EV isolation.

We propose that the use of Opti-MEM for sample collection, should be avoided (or at least carefully controlled) for studies of EVs and conditioned media transfer in myogenic cultures (and potentially in other insulin-sensitive biological systems). Similarly, polymer precipitation methods of EV isolation should both be avoided due to the risk of introducing such confounding factors.

