## Supplementary Information for "Extracellular vesicle-mediated promotion of myogenic differentiation is dependent on dose, collection media composition, and isolation method"

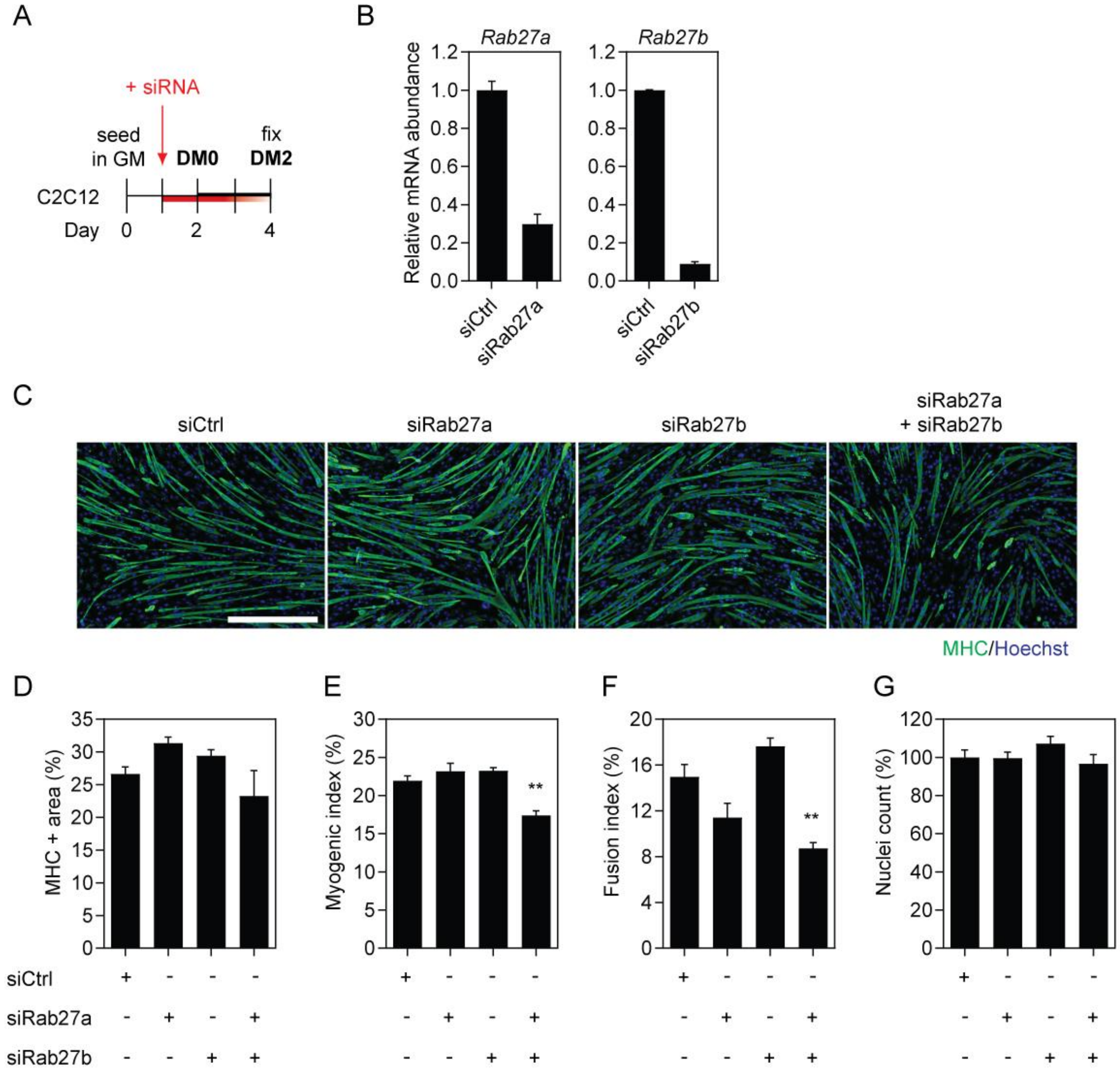

**Figure S1**

**Combined siRNA-mediated knock down of *Rab27a* and *Rab27b* exosome biogenesis factors reduces myogenic differentiation.**

(**A**) C2C12 cells were cultured in GM for two days and then switched to DM for two days. Cultures were treated with siRNAs targeted to two exosome biogenesis factors, *Rab27a* and *Rab27b*, separately or in combination, at a final concentration of 100 nM one day before switching to DM. Treatment with a non-targeting siRNA was included as a negative control. (**B**) *Rab27a* and *Rab27b* mRNA levels were determined by RT-qPCR, normalised to the *Rplp0* reference gene. The data were scaled such that the mean of the control group was returned to a value of 1 ($n$=2). (**C**) Myogenic differentiation was assessed by MHC IF, and quantified by (**D**) measuring the MHC+ area, and by calculating the (**E**) myogenic and (**F**) fusion indices. (**G**) The total number of nuclei per representative field of view are shown as a percentage relative to the control group. Cultures treated with a non-targeting siRNA pool were included as negative controls. All microscopy images were taken at 10× magnification. Scale bar represents 400 µm. Values are mean + SEM ($n$=4). Statistical significance was determined by one-way ANOVA with Bonferroni *post hoc* test, $^{**}P<0.01$.

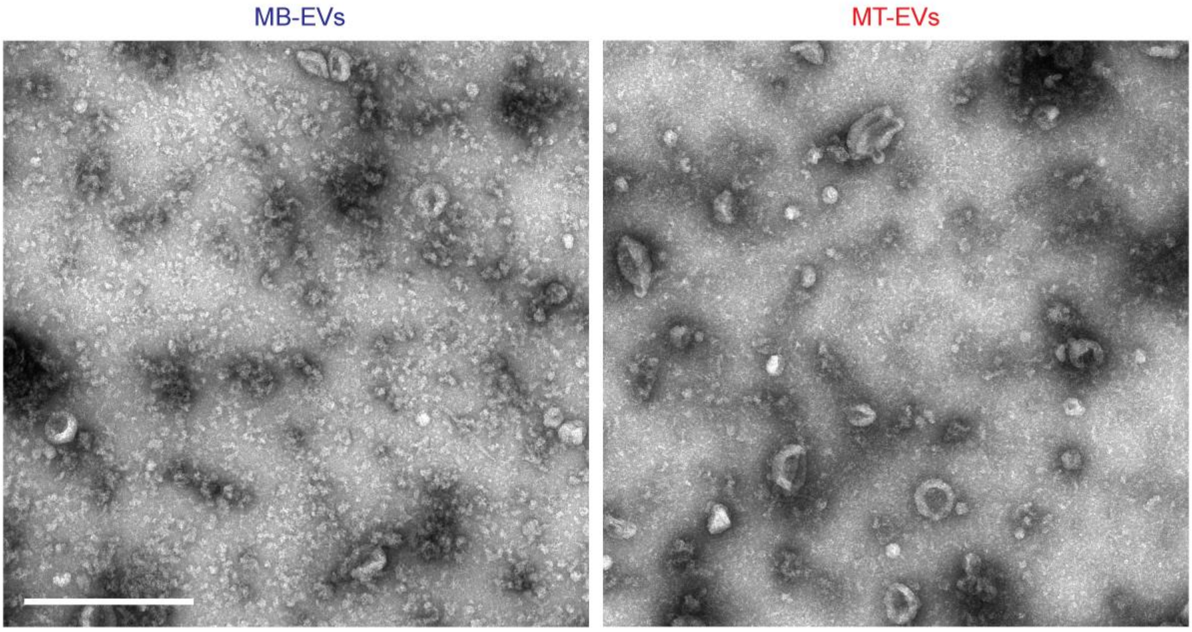


**Figure S2**

**Analysis of EV preparations by transmission electron microscopy.**

EVs were collected from myoblasts (MBs) or myotubes (MTs) as described in **Figure 3A** and preparations were analysed by transmission electron microscopy. Vesicles with 'cup-shaped' morphology characteristic of exosomes are observed. Scale bars represent 500 nm.

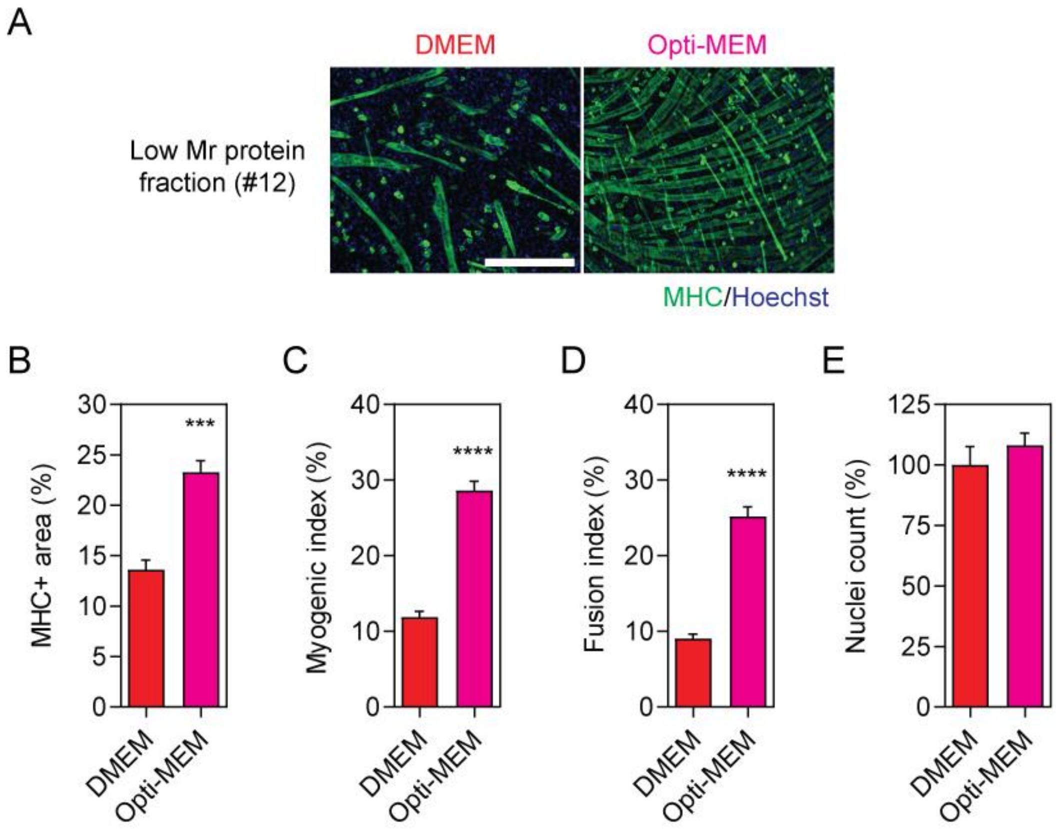


**Figure S3**

**Low molecular weight protein obtained from unconditioned Opti-MEM media is highly pro-myogenic.**

Fresh Opti-MEM and DMEM were fractionated by UF-SEC and protein fraction #12 collected as shown in **Figure 5B**. Recipient C2C12 cultures were treated with 1 µg/ml of purified protein fraction at the time of switching to DM. (**A**) Myogenic differentiation was assessed by MHC IF at DM2 and quantified by measuring (**B**) MHC+ area, (**C**) myogenic index, and (**D**) fusion index. (**E**) The total number of nuclei per representative field of view are shown as a percentage relative to the control group. All microscopy images were taken at 10× magnification. Scale bar represents 400 µm. All values are mean + SEM (*n*=4). Statistical significance was determined by a Student's *t* test, ***$P$<0.001, ****$P$<0.0001.

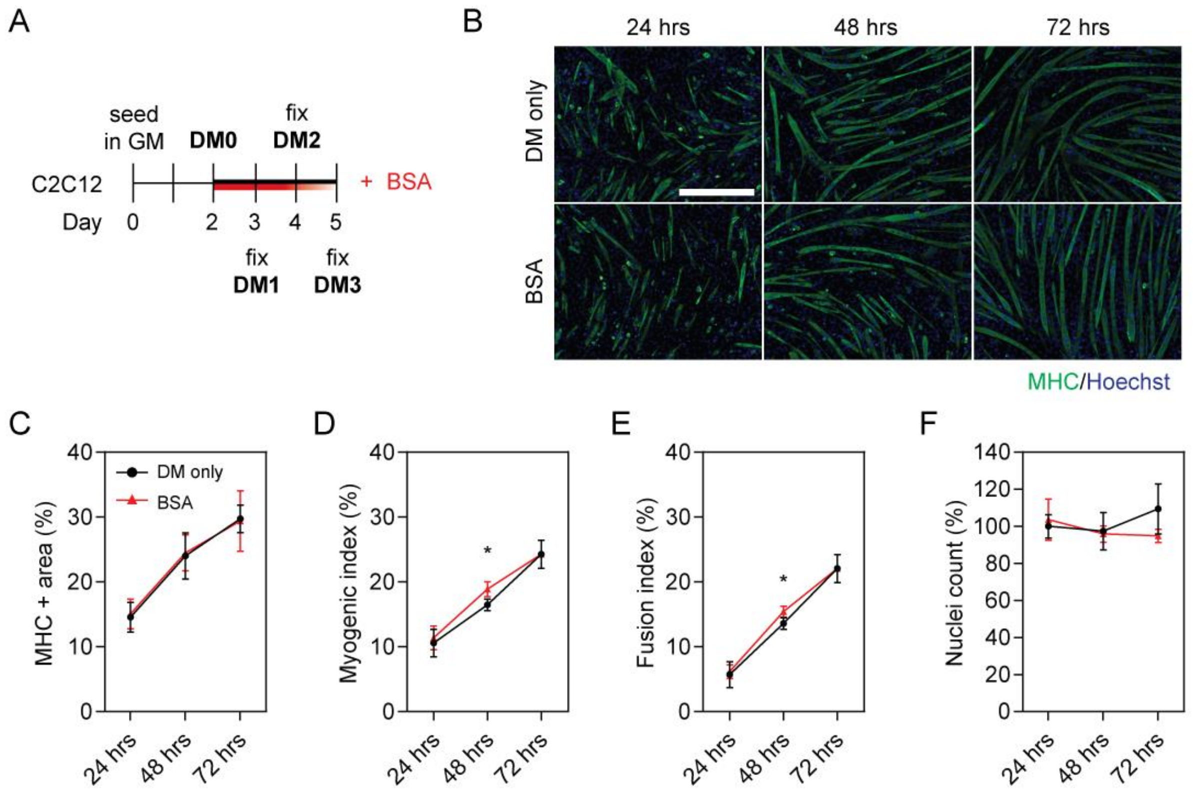


**Figure S4**

**A non-specific increase in extracellular protein does not enhance myogenic differentiation.**

(**A**) C2C12 myoblasts were grown in GM for two days, and treated with 5 μg/ml of bovine serum albumin (BSA) at the time of switching to DM. (**B**) Myogenic differentiation was assessed by MHC IF at DM1, DM2 and DM3, and quantified by measuring (**C**) MHC+ area, (**D**) myogenic index, and (**E**) fusion index. Untreated (DM only) cultures were included as negative controls. (**F**) The total number of nuclei per representative field of view are shown as a percentage relative to the control group. All microscopy images were taken at 10× magnification. Scale bar represents 400 μm. All values are mean ± SEM ($n$=4). Statistical significance was determined by Student's $t$ test, *$P<0.05$.

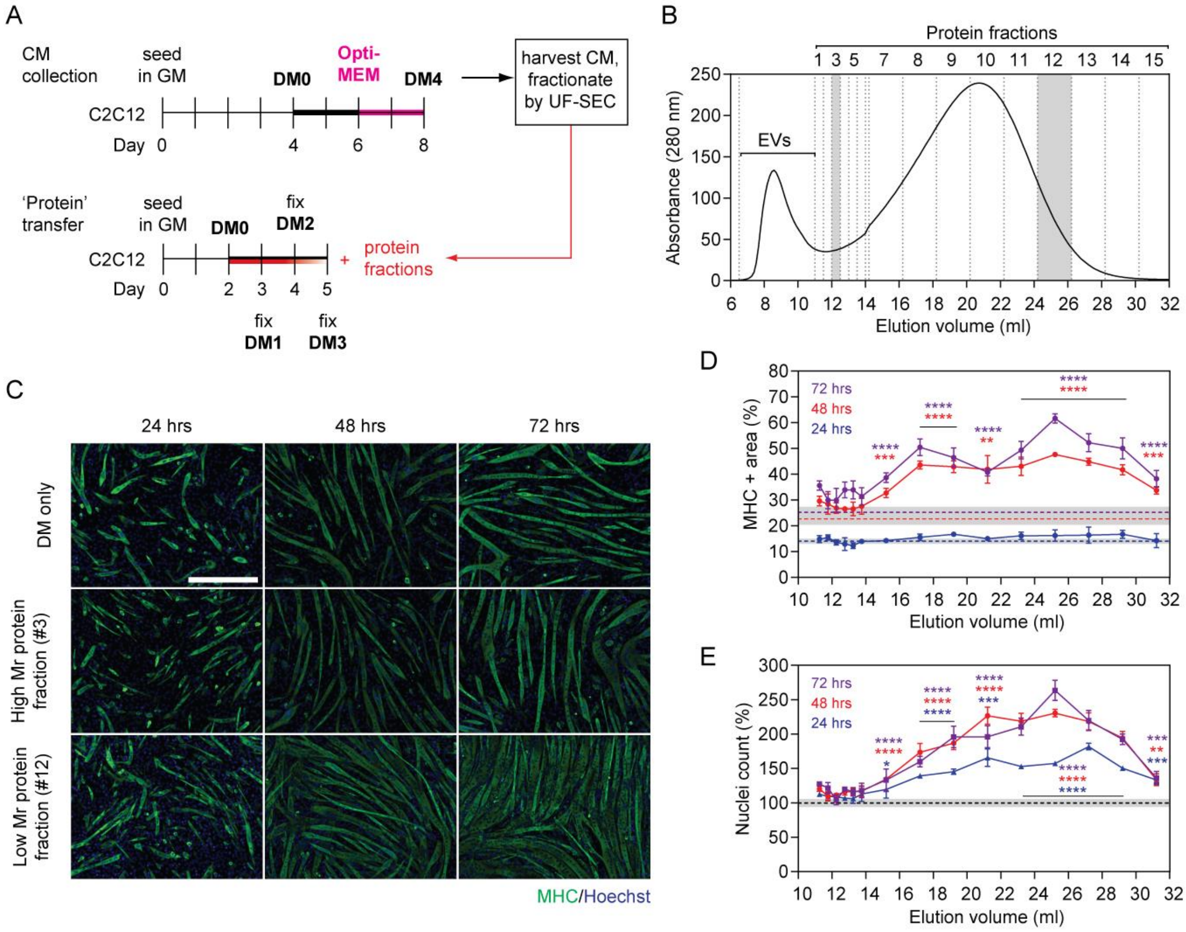
A
CM collection
seed in GM
Opti-MEM
DM0
DM4
harvest CM, fractionate by UF-SEC
C2C12
Day 0 4 6 8
'Protein' transfer
seed in GM
DM0
fix DM2
C2C12
+ protein fractions
Day 0 2 3 4 5
fix DM1
fix DM3
B
Protein fractions
1 3 5 7 8 9 10 11 12 13 14 15
EVs
Absorbance (280 nm)
Elution volume (ml)
C
24 hrs
48 hrs
72 hrs
DM only
High Mr protein fraction (#3)
Low Mr protein fraction (#12)
MHC/Hoechst
D
72 hrs
48 hrs
24 hrs
MHC + area (%)
Elution volume (ml)
E
72 hrs
48 hrs
24 hrs
Nuclei count (%)
Elution volume (ml)

**Figure S5**

**Low molecular weight myotube-derived secreted protein enhances myogenic differentiation when purified from Opti-MEM collection media.**

(**A**) C2C12 myotube-derived secreted protein fractions were isolated by UF-SEC from CM collected in Opti-MEM. Recipient cultures were treated with 1 μg/ml of purified extracellular protein from each fraction at the time of switching to DM. (**B**) The non-vesicular-associated protein eluates from the SEC column were measured by UV spectrophotometry absorbance at 280 nm. Two fractions (#3 and #12) were selected to represent high and low Mr extracellular protein fractions, respectively (indicated by grey shading). (**C**) Myogenic differentiation in recipient cultures was assessed by MHC IF at DM1 (24 hours), DM2 (48 hours), and DM3 (72 hours). Representative images from the high and low Mr extracellular protein fractions are shown from each time point. (**D**) Myogenic differentiation was quantified by measuring the MHC+ area. The means of corresponding DM only control groups from each time point are shown ± SEM (grey shaded area). (**E**) The total number of nuclei per representative field of view are shown as a percentage relative to the control group (data were scaled such that the mean of the control group was returned to a value of 100%). Untreated (DM only) cultures were included as negative controls. All microscopy images were taken at 10× magnification. Scale bar represents 400 μm. Values are mean ± SEM ($n$=4). Statistical significance was determined by a Student's $t$-test, $^{**}P<0.01$, $^{***}P<0.001$, and $^{****}P<0.0001$ (for ease of interpretation test results are only shown for fraction #7-15). The colour of the significance indicators corresponds to the respective comparison relative to the DM only control at each time point (i.e. blue for the 24 hour time point, red for 48 hours, and purple for 72 hours).

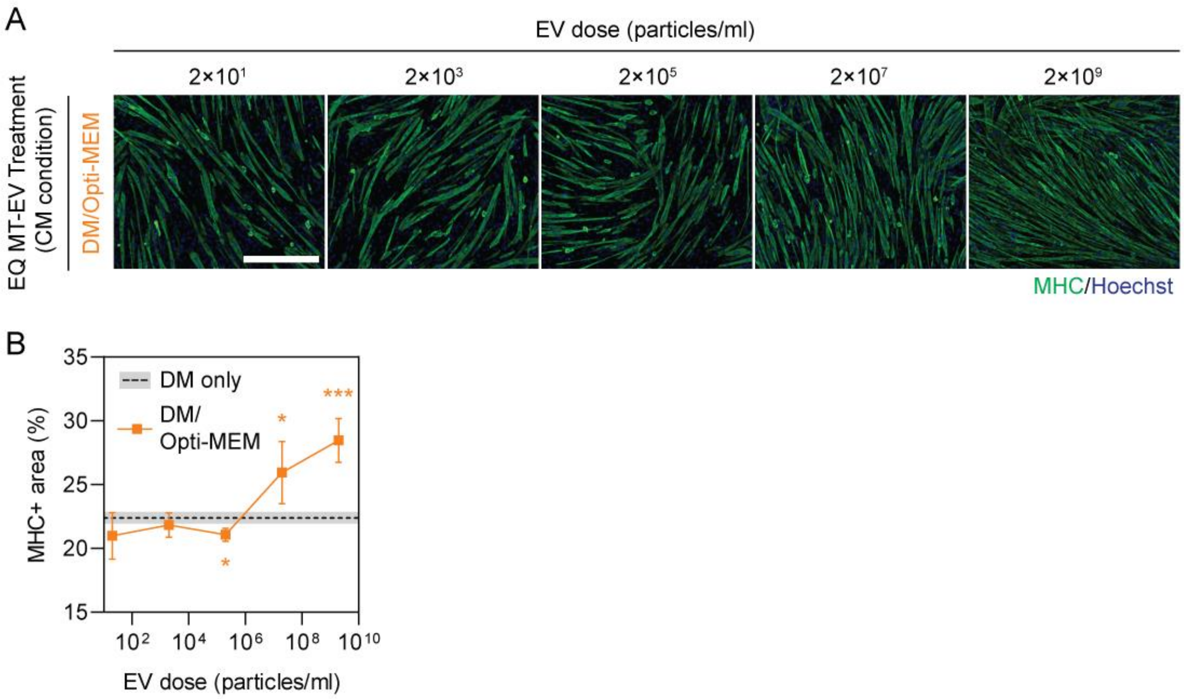
A
EV dose (particles/ml)
2×10¹
2×10³
2×10⁵
2×10⁷
2×10⁹
EQ MT-EV Treatment (CM condition)
DM/Opti-MEM
MHC/Hoechst
B
DM only
DM/ Opti-MEM
MHC+ area (%)
EV dose (particles/ml)

**Figure S6**

**EQ MT-EVs obtained from Opti-MEM collection media exhibit pro-myogenic effects at high doses.**

EVs were isolated by ExoQuick polymer precipitation (EQ MT-EVs) from C2C12 myotubes and collected in Opti-MEM isolation media. Myoblasts were grown in GM for four days, DM for two days, and isolation medium (Opti-MEM) for a further two days. Recipient myoblast cultures were treated with a range of EQ MT-EV doses. (**A**) Myogenic differentiation was assessed by MHC IF at DM2 and quantified by (**B**) measuring the MHC+ area. All microscopy images were taken at 10× magnification. Scale bar represents 400 µm. All values are mean ± SEM ($n$=4). Untreated (DM only) cultures were included as negative controls. Statistical significance was determined by Student's $t$-test, *$P<0.05$, ***$P<0.001$.

| ID | Target sequence (5' to 3') |
|---|---|
| siRab27a | CGGAUGGAGAUUACGAUUA |
| | CAGGAGAGGUUUCGUAGCU |
| | GUACAGAGCCAAUGGGCCA |
| | GGGCAUUGAUUUCAGGGAA |
| siRab27b | GCAGAUUAGAAGCUAGUAU |
| | GGGAAUAGAUUUUCGGGAA |
| | GGAAGUCAAUGAACGGCAA |
| | GCAGAGUAGUCAUAGUGUU |
| siCtrl | UGGUUUACAUGUCGACUAA |
| | UGGUUUACAUGUUGUGUGA |
| | UGGUUUACAUGUUUUCUGA |
| | UGGUUUACAUGUUUUCCUA |

**Table S1**

**Target sequences for SMARTpool siRNAs used in this study.**

| Target protein | Host | Product ID | Manufacturer | Dilution |
|---|---|---|---|---|
| **Western Blot** | | | | |
| **Primary Abs** | | | | |
| Anti-PDCD6IP (Alix) | Mouse | ab117600 | Abcam | 1:1,000 |
| Anti-TSG101 | Mouse | 612696 | BD Biosciences | 1:1,000 |
| Anti-CANX (Calnexin) | Rabbit | ab22595 | Abcam | 1:1,000 |
| Anti-CD9 | Rabbit | ab92726 | Abcam | 1:2,000 |
| **Secondary Abs** | | | | |
| Anti-mouse IgG, HRP-linked | Horse | 7076 | Cell Signaling Technology | 1:5,000 |
| **Immunofluorescence** | | | | |
| **Primary Abs** | | | | |
| Anti-MHC | Mouse | MF 20 | Abcam | 1:20 |
| **Secondary Abs** | | | | |
| Anti-rat IgG Alexa Fluor 488 | Goat | ab150157 | Abcam | 1:500 |

**Table S2**

**Antibodies used in this study.**

MF 20 primary monoclonal antibody, developed by Fischman, D.A. at Weill Cornell Medical College, was obtained from the Developmental Studies Hybridoma Bank, created by the NICHD of the NIH and maintained at The University of Iowa, Department of Biology, Iowa City, IA 52242, USA. WB, western blot, IF, immunofluorescence.

| ID | Sequence (5' to 3') |
|---|---|
| **qRab27a-Fwd** | CGACCTGACAAATGAGCAAAG |
| **qRab27a-Rev** | CCTCTTTCACTGCCCTCTG |
| **qRab27b-Fwd** | CAGACCTGCCAGACCAAAG |
| **qRab27b-Rev** | AGCGTTTCCACTGACTTCTC |
| **qRplp0-Fwd** | AAGCAAAGGAAGAGTCGGAG |
| **qRplp0-Rev** | CCAGACCGGAGTTTTAAGAGAAG |

**Table S3**

**RT-qPCR primer sequences used in this study.**

| Target | Product ID | Detection Channel |
|---|---|---|
| **mmu-miR-1a-3p** | 002222 | FAM |
| **mmu-miR-133a-3p** | 002246 | FAM |
| **mmu-miR-206-3p** | 000510 | FAM |
| **cel-miR-39** | 000200 | FAM |

**Table S4**

**miRNA small RNA TaqMan RT-qPCR assays used in this study.**

All products were purchased from Thermo Fisher Scientific.

| ID | Sequence (5' to 3') |
|---|---|
| **mmu-miR-1a-3p** | UGGAAUGUAAAGAAGUAUGUAU |
| **mmu-miR-133a-3p** | UUUGGUCCCCUUCAACCAGCUG |
| **mmu-miR-206-3p** | UGGAAUGUAAGGAAGUGUGUGG |
| **cel-miR-39** | UCACCGGGUGUAAAUCAGCUUG |

**Table S5**

**List of oligonucleotides used in this study.**

Oligonucleotides were purchased from IDT (Leuven, Belgium).
